# The assembly of stress granule-like foci during foot-and-mouth disease virus infection is uncoupled from activation of cellular intrinsic antiviral signalling

**DOI:** 10.1101/2025.10.30.685519

**Authors:** Mariana Marques, Zhaozhi Sun, Amina Yasmin, J. Paul Taylor, Alessia Ruggieri, Tobias J. Tuthill, Nicolas Locker

**Affiliations:** The Pirbright Institute, GU24 0NF Pirbright, UK; Heidelberg University, Medical Faculty Heidelberg, Department of Infectious Diseases, Molecular Virology, Centre for Integrative Infectious Disease Research, 69120 Heidelberg, Germany; Department of Cell and Molecular Biology, St. Jude Children’s Research Hospital, Memphis, TN 38105, USA

## Abstract

Foot-and-mouth disease virus (FMDV) is highly contagious among cloven-hoofed animals and poses a major threat to the livestock industry worldwide. A fundamental gap in knowledge for high consequence viruses such as FMDV is understanding how the virus evolved to evade cellular antiviral responses. FMDV belongs to the *Picornaviridae*, a family of positive-sense single-stranded RNA viruses. The detection of viral double-stranded viral RNA intermediates during infection can trigger both the assembly of cytoplasmic stress granules (SGs) and the activation of the RIG-I-like receptors (RLR)-mediated innate immune response (IIR). FMDV has been proposed to antagonize these mechanisms, suggesting that both can limit viral replication. In this study, we investigate the dynamic and importance of SG assembly for IIR activation upon dsRNA stimulation or FMDV replication in porcine epithelial kidney cells.

First, we show that the formation of SG following a challenge with poly(I:C), a viral dsRNA mimic, does not modulate the activation of IIR. Our data further reveal transient assembly of SG during FMDV infection followed by virus-induced cleavage of G3BP1, a core SG protein. While SG assembly does not impact viral replication or antiviral response activation, we demonstrate that preventing their disassembly negatively impacts FMDV replication. Overall, our data suggests that while SGs assembly is uncoupled from IIR activation, manipulating their disassembly limits replication and might serve as a potential therapeutic target to prevent FMDV infection.

**Importance:** Biomolecular condensates, including stress granules, are key regulatory compartments that control fundamental cellular processes. By condensing RNAs and proteins, these structures enable cells to rapidly adapt to stress, such as viral infection. Different biomolecular condensates with specific dynamics or compositions have been described during infection, and many viruses are known to disrupt or highjack their components. Moreover, the cell-autonomous innate immune response is proposed to be regulated by biomolecular condensates. However, the molecular mechanisms underpinning these functions remain unclear and controversial. Here we investigated the interplay between stress granules and the innate response upon the stimulation with a viral dsRNA mimic or foot-and-mouth disease virus infection. We demonstrate that infection triggers the formation of stress granules independently from activation of innate signalling pathways. We also show that stress granules persistence attenuates viral replication posing them as part of the cell’s response that viruses must overcome or subvert to replicate.

## Introduction

Foot-and-mouth disease virus (FMDV) is the aetiological agent of foot-and-mouth disease (FMD), a highly contagious diseases that infects a broad range of wild and domestic cloven-hoofed animals (1). FMD is endemic in Africa and Asia, where it is continuously controlled by vaccination, yet outbreaks in FMD-free regions, as recently in Europe, require strict restrictions (2). Causing severe pain and distress in the infected animals, FMD seriously affect livestock and its industry worldwide, threatening the international trade in animals and animal products with severe economic implications (3).

FMDV belongs to the family *Picornaviridae* of non-enveloped viruses with single-stranded positive-sense RNA genome, which includes a long 5′-untranslated region (5′UTR), a large open reading frame (ORF), and a short 3′UTR. Translation begins at the IRES element in the 5’UTR to produce a polyprotein encoding for eight non-structural proteins that regulate RNA replication, protein folding and virus assembly, and four structural proteins, constituting the capsid (4).

During RNA virus infection, the detection of double-stranded RNA (dsRNA) replication intermediates or deregulation of cellular homeostasis activates the integrated stress response (ISR), which results in an adaptive transcriptional and translational rewiring to limit viral protein synthesis and promote cellular recovery (5). This process is mediated by the phosphorylation of the eukaryotic translation initiation factor 2α (eIF2α) and involves polysome disassembly and the consequent accumulation of stalled mRNAs, which serve as scaffold to nucleate RNA-binding proteins (RBPs), such as Ras-GTPase activating protein-binding protein 1 (G3BP1). This triggers the recruitment and accumulation of proteins into ribonucleoprotein complexes that ultimately mature into cytoplasmic membrane-less condensates, named stress granules (SGs), by liquid-liquid phase separation (6–10). SGs are highly heterogeneous and dynamic, rapidly assembling in response to stress and dissolving upon its resolution to resume the translation of stalled mRNAs (11). Several studies have identified and characterized SG-like or related condensates during infection, including paracrine granules and RNAse-L bodies (RLBs) that differ from canonical SGs in terms of composition and function (5, 12, 13). These observations highlight the diverse nature of stress-induced biocondensates, and the importance of context and composition for their biological function.

The exact function of SGs and SG-like condensates during viral infection and their interplay with the cell intrinsic antiviral innate immune response (IIR) is unclear. The detection of dsRNA intermediates triggers an IIR mediated by cytoplasmic RIG-I-like receptors (RLR) (14), relaying a signalling cascade mediated by mitochondrial antiviral signalling (MAVS) to trigger an interferon (IFN)-mediated transcriptional induction of antiviral genes (15). SGs have been proposed to function as either antiviral platforms or signalling scaffolds for RLRs or other antiviral effectors (16–22). In turn, numerous viruses have evolved strategies to antagonize SG formation or promote their disassembly (5). SGs have also been described as “shock absorbers” that dampen antiviral signalling by sequestering antiviral factors and preventing or delaying their activation to protect cells from immune-mediated apoptosis (23). Alternatively, SGs could be simple by-products of viral infections and independent from antiviral immune responses (12, 18, 24).

FMDV has been previously shown to interfere with host translation and SG components to evade these mechanisms. Both FMDV proteases, Lpro and 3Cpro, are associated with cleavage of G3BP1 to antagonize SGs, yet whether SGs assemble during infection or their impact on IFN signalling remains unclear (25, 26).

Herein, we investigated the significance of SGs during RNA virus infection in porcine epithelial kidney cells (PK-15) and explored their relevance for the activation of the intrinsic IIR. First, we show that the induction of IIR signalling and the assembly of G3BP1-foci are uncoupled following the stimulation with poly(I:C), a synthetic viral dsRNA mimic. We also demonstrate that stimulation of the RIG-I/MAVS pathway is not sufficient to induce the formation of G3BP1-foci. Furthermore, our data show that FMDV infection promotes SGs assembly followed by their rapid disassembly and cytopathic effect. Inhibiting G3BP1 condensation and foci assembly does not impact FMDV replication, neither modulates the IIR against a mutated FMDV that lacks the ability to evade IFN signalling. Lastly, our data indicate that delaying the disassembly of FMDV-induced G3BP1-foci during infection reduces FMDV replication. Overall, this sheds light on the interplay between of biocondensates dynamics and IIR signalling during infection.

## Results

### The IIR is induced following stimulation with viral dsRNA mimics in PK-15 cells

PK-15 cells are porcine kidney epithelial cells commonly used to study FMDV infection. To ensure appropriate monitoring of ISR and IIR induction, we established that this cell line is responsive to synthetic viral dsRNA mimics, such as 3p-hpRNA and poly(I:C). 3p-hpRNA is a specific RIG-I ligand, consisting of a short RNA hairpin structure with an uncapped 5’ triphosphate extremity (27), whereas poly(I:C) is recognized by several cytosolic receptors, including RIG-I-like receptors (RLR) (28), protein kinase R (PKR) (12) and oligoadenylate synthetase-like (OASL) (29). To characterize the antiviral response initiated by 3p-hpRNA and poly(I:C) in PK-15, cells were transfected for 6 hours with increasing amounts of each viral mimic prior to measuring the levels of IIR and ISR factors using RT-qPCR or western blot (Fig. **1A-B**).

**Figure 1.**
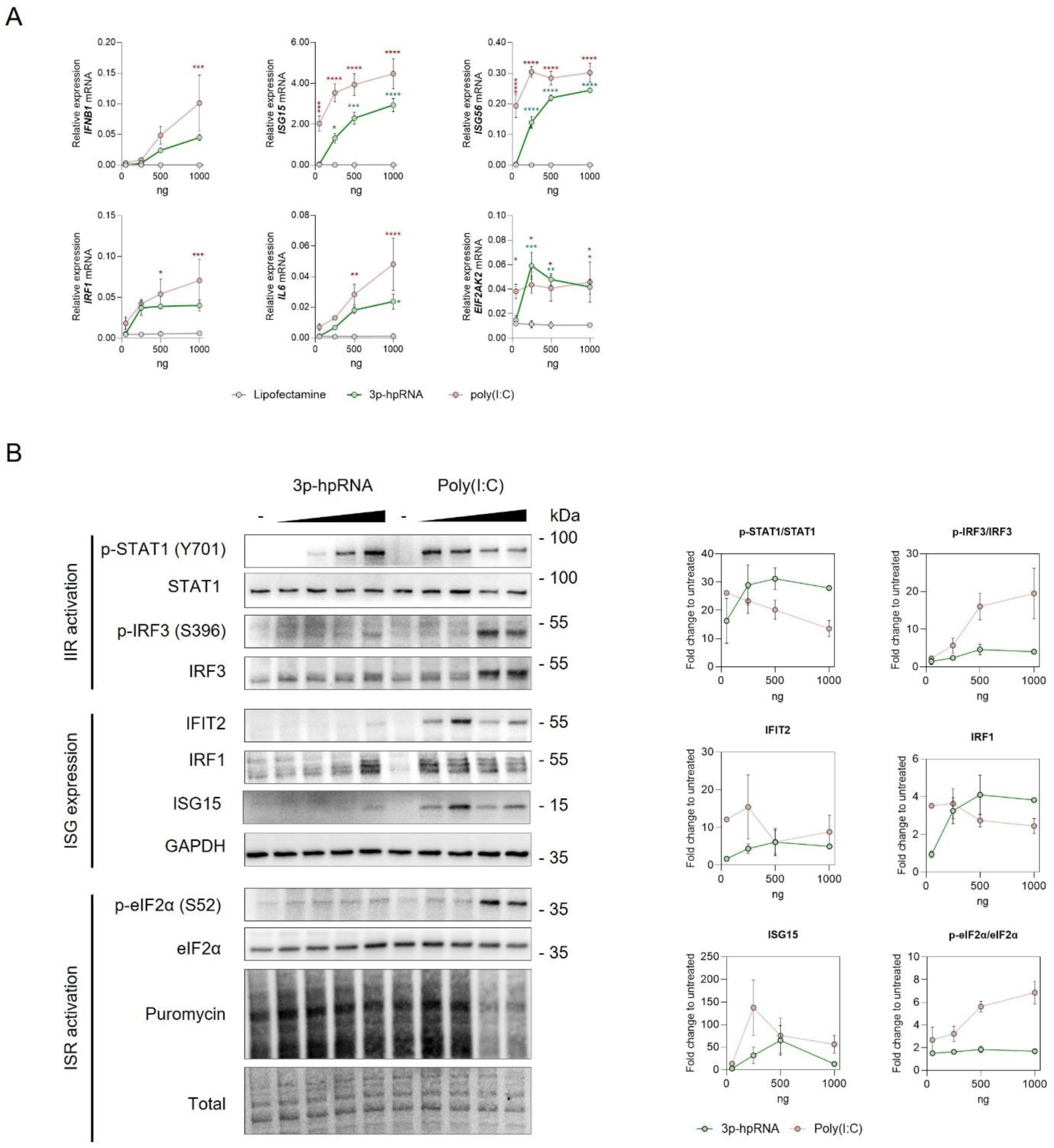
IIR is robustly induced following PKR and RIG-I stimulation in PK-15 cells. PK-15 cells were transfected with increasing amounts of 3p-hpRNA or poly(I:C) for 6 hours. Transfection with lipofectamine was used as a control. (A) Expression levels of *IFNB1* and ISGs were analysed by RT-qPCR with results shown as mean ± SEM, n=3, normalised to *GAPDH* mRNA. *p<0.05, **p<0.1, *** p<0.001, ****p<0.0001; using two-way ANOVA with Šidák’s multiple comparison between 3p-hpRNA (in green) or poly(I:C) (in red) and lipofectamine. (B) Representative western blot shown and quantification from at least three independent experiments for markers of interferon signalling activation (STAT1 and IRF3 phosphorylation levels), interferon-stimulated genes expression (IFIT2, IRF1 and ISG15), and integrated stress response activation (phosphorylation of eIF2α and puromycin levels). Molecular weights are indicated on the right. Band intensities were normalised to that of total protein or GAPDH. Data represents mean ± SEM of three biological replicates.

As expected, increasing amounts of 3p-hpRNA resulted in a dose-dependent induction of *IFNB1* and ISGs expression, including *ISG15*, *ISG56*, *IRF1*, *IL6*, *and EIF2AK2*, as seen by qPCR (Fig. **1A**). Similarly, we detected higher levels of IRF3 and STAT1 phosphorylation and ISGs, such as IFIT2, IRF1, ISG15 using immunoblotting (Fig. **1B**). Transfection with increasing amounts of poly(I:C) promoted a stronger *IFNB1* and ISGs expression (Fig. **1A**), accompanied by higher levels of IRF3 phosphorylation (Fig. **1B**), but a plateau in the phosphorylation of STAT1 and ISGs synthesis (IFIT2, IRF1, ISG15) (Fig. **1B**). These observations might be explained by the activation of the ISR and resulting decrease in protein synthesis following poly(I:C) but not 3p-hpRNA stimulation, detected by the phosphorylation of eIF2α and puromycin levels, respectively (Fig. **1B**). In summary, these data support the use of viral mimics to induce antiviral response in our model system.

### Stimulation with poly(I:C), but not 3p-hpRNA, induces the assembly of G3BP1-foci

Next, we set out to dissect the assembly of SGs in PK-15 cells following a 6-hour treatment with increasing amounts of poly(I:C) or 3p-hpRNA. Sodium arsenite was used as a positive control for the assembly of canonical eIF2α-dependent SGs (30). Cells were stained using the SG markers G3BP1 and eIF4GI (Fig. **2A-B**). In untreated cells, G3BP1 and eIF4GI remained diffuse in the cytoplasm while arsenite treatment resulted in accumulation of G3BP1/eIF4Gs into foci in around 80% of the cells, reflecting SG assembly, with an average number of 28.9±6.2 foci per cell with an average size of 0.61±0.08 μm^2^. Treatment with increasing amount of poly(I:C) resulted in the dose-dependent induction of foci that appeared less numerous and smaller in size when compared to canonical arsenite-induced SGs (Fig. **2A-B**). In contrast, 3p-hpRNA did not induced the assembly of G3BP1-foci at any of the concentrations tested (Fig. **2A-B** and **S1A**). These data suggest that the assembly of G3BP1-foci is triggered independently of the RIG-I/MAVS-mediated response and therefore relies on the poly(I:C) stimulation of other receptors.

**Figure 2.**
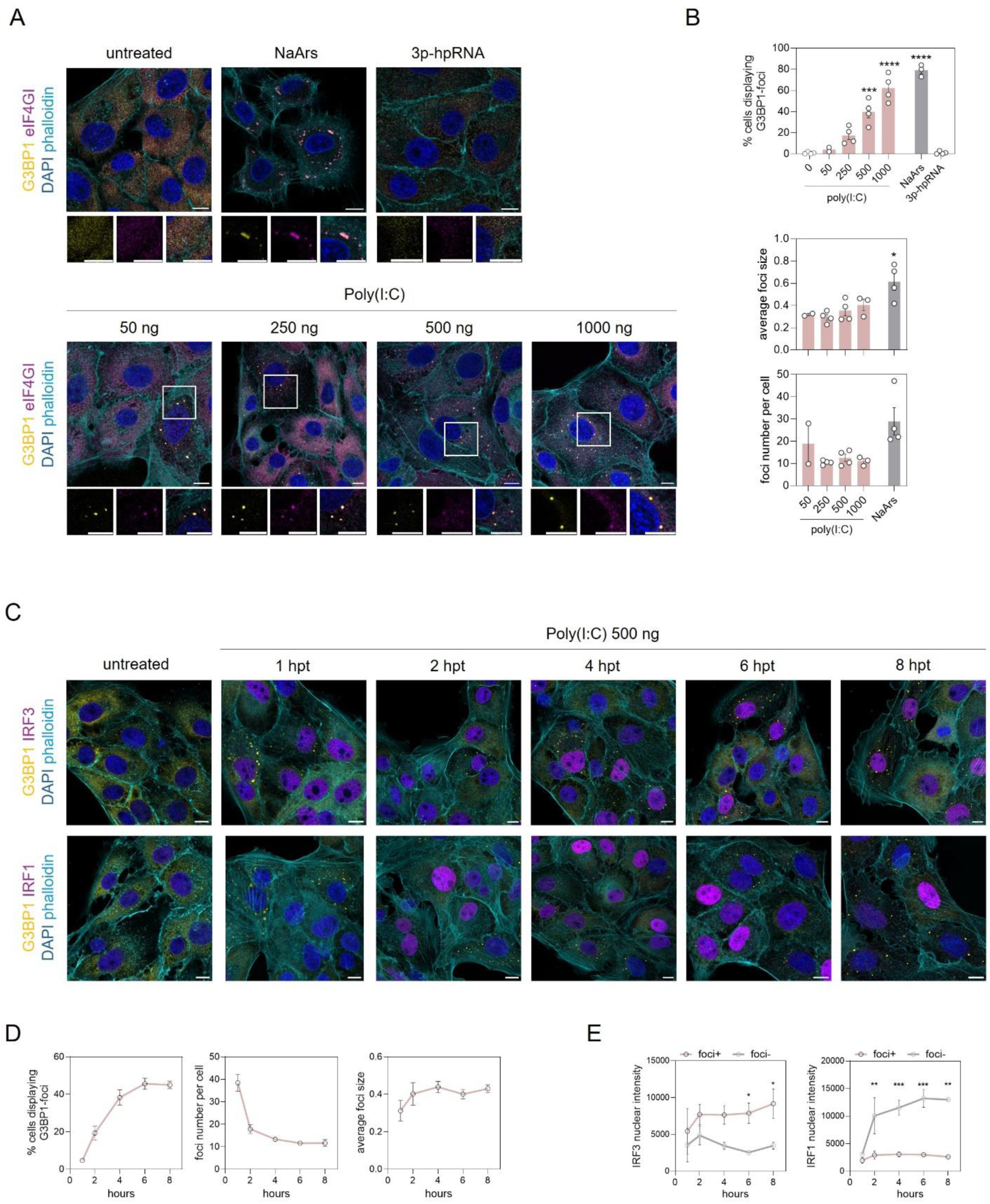
Poly(I:C), but not 3p-hpRNA, induces G3BP1-foci formation and IIR activation at similar times. (A) Representative confocal images and (B) quantified data of PK-15 cells stimulated with 3p-hpRNA or increasing concentrations of poly(I:C) for 6 hours. Sodium arsenite treatment (1 mM, 1 hour) was used as positive control. Cells were analysed by immunofluorescence for the SG markers G3BP1 (gold) and eIF4GI (magenta), and a F-actin marker phalloidin (cyan). Nuclei were stained with DAPI. Scale bars represent 10 μm. The percentage of cells displaying G3BP1-foci was quantified by manual counting of at least 100 cells per replicate. The average size of foci per cell and the number of foci per cell were quantified using the G3BP1 channel and ‘Analyse Particles’ plugin from ImageJ. Results shown as mean ± SEM of at least three independent biological replicates, representing the average of at least 30 cells per condition. *p<0.05, *** p<0.001, ****p<0.0001; using ordinary one-way ANOVA with Šidák’s multiple comparison post-test to 50ng poly(I:C). (C) Confocal images of PK-15 cells stimulated with 1 μg/ml poly(I:C) up to 8 hours post stimulation. Cells were analysed by immunofluorescence for the SG marker G3BP1 (gold), innate immune response activation markers IRF3 or IRF1 (magenta), and F-actin marker phalloidin (cyan). Nuclei were stained with DAPI. Scale bars represent 10 μm. (D) Quantification of the percentage of cells displaying G3BP1-foci, number of foci per cell, average number and size of foci per cell. Data represents mean ± SEM of three biological replicates. (E) Quantification of IRF1 or IRF3 nuclear intensities in cells positive or negative for SGs was done using ImageJ. *p<0.05, **p<0.1, *** p<0.001; using two-way ANOVA with Šidák’s multiple comparison between cells with (foci+) and without (foci-) G3BP1 condensates at each time point.

To address the dynamics of ISR and IIR activation following poly(I:C) stimulation, we conducted a time-course experiment from 1 to 6 hours post-transfection (hpt) and monitored both the assembly of G3BP1-foci and IIR activation using immunofluorescence assays, labelling for G3BP1 as SG/ISR marker, or IRF3 and IRF1 as marker of IIR signalling (Fig. **2C-E**). Upon RIG-I/MAVS signalling, the activation of IRF3 results in its intrinsic nuclear translocation, followed by an interferon-mediated expression of ISG in both autocrine and paracrine ways. The nuclear localization of IRF1 was used as measure of ISG expression, as this feature is lost upon inhibiting interferon-mediated signalling with ruxolitinib (**Fig. S1B**).

Following poly(I:C) treatment, G3BP1-foci assembled in around 5% of the cells from 1 hpt, reaching 45% of the cells at 6 hpt, which was maintained until 8 hpt. The number of foci per cell was higher at 1 hpt (38.5±3.8) compared to the later time points (17.9±1.9, 13.4±0.7, 11.6±0.8 and 11.6±1.6 for 2, 4, 6 and 8 hpt, respectively), whereas their size at 1 hpt was smaller (0.31±0.05 μm^2^) compared to later times (0.40±0.06, 0.44±0.03, 0.40±0.02 and 0.43±0.02 μm^2^) (Fig. **2D**). This may reflect the proposed multistep process for SGs assembly whereby small aggregates coalesce before fusing into larger condensates (31).

Our analysis revealed that the activation of the RIG-I/MAVS/IRF3 pathway and the assembly of G3BP1-foci occur within the same cell, as reflected by the intensity levels of nuclear IRF3 in cells with and without foci (Fig. **2E**). Interestingly, the nuclear localization of IRF1 was not detected in cells assembling G3BP1-foci upon poly(I:C) transfection (Fig. **2E**), suggesting these cells are unable to produce antiviral factors to the same extent. Overall, this suggests that IIR and ISR activation occur at a similar time, raising the question of whether one drives the other, and suggests that ISR activation might impair the IFN-mediated response.

### Inhibiting G3BP1 condensation blocks poly(I:C)-induced foci, but does not revert IIR signalling

To establish whether the ability to assemble SGs would impact IIR induction, we leveraged a pharmacological inhibitor of SG assembly, the G3BP condensation inhibitor b (G3Ib), which binds the G3BP1 domain essential for the conformational change driving SG assembly (32). This allows to discriminate between the role of SGs from that of their individual components, presenting an advantage over G3BP1/2 knocked out genetic models, which does not discriminate between loss of G3BP1/2 SG-dependent functions and SG-independent functions, such as modulating the innate immune response or RNA metabolism (19, 21).

We first tested the ability of G3Ib to prevent the assembly of poly(I:C)-induced SGs and their relevance for IIR activation, using its inactive enantiomer G3Ib’ as a control. To visualize G3BP1-foci assembly and IIR activation, cells were labelled for G3BP1 and IRF3 or IRF1, respectively (Fig. **3A**). Treating cells with G3Ib resulted in impaired foci assembly following poly(I:C) compared to untreated cells, with reduction in percentage of cells with G3BP1-foci from 56.3 to 12%, while the inactive drug G3Ib’ had no impact (Fig. **3B**). The number of G3BP1-foci per cell and average size were also reduced in cells treated with G3Ib (2.08±0.16 and 0.10±0.01 μm^2^) compared to poly(I:C) treatment alone (12.41±0.79 and 0.32±0.01 μm^2^), while G3Ib’ had no significant effect (12.82±1.34 and 0.34±0.01 μm^2^) (Fig. **3B**). These data further support the use of G3Ib as a tool to prevent the assembly of poly(I:C)-induced G3BP1-foci and characterise their functions during IIR activation.

**Figure 3.**
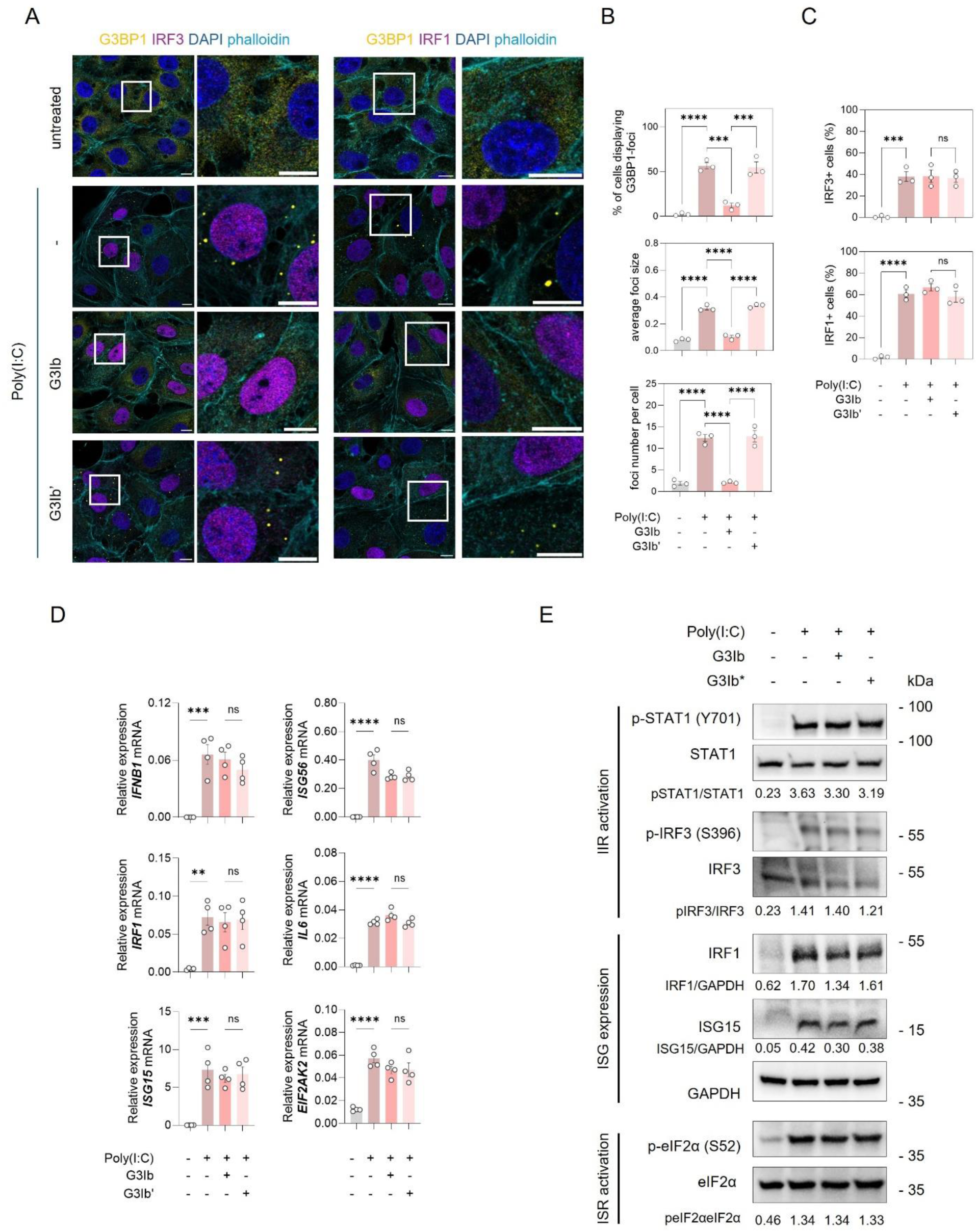
Inhibiting G3BP1 condensation blocks poly(I:C)-induced foci. (A) Representative confocal images and (B) quantified data of PK-15 cells stimulated with poly(I:C) for 6 hours following treatment with G3BP1 condensation inhibitor (G3Ib) or its enactive enantiomer (G3Ib’). Cells were analysed by immunofluorescence for the SG markers G3BP1 (gold), innate immune response activation markers IRF3 or IRF1 (magenta), and a F-actin marker phalloidin (cyan). Nuclei were stained with DAPI. Scale bars represent 10 μm. The percentage of cells displaying G3BP1-foci was quantified by manual counting of at least 100 cells per replicate. The average size of foci per cell and the number of foci per cell were quantified using the G3BP1 channel and ‘Analyse Particles’ plugin from ImageJ. Results shown as mean ± SEM of at least three independent biological replicates, representing the average of at least 30 cells per condition. (C) Quantification of IRF1 or IRF3 nuclear intensities in cells positive or negative for SGs was done using ImageJ. (D-E) Expression levels of IFNβ and ISGs were analysed by RT-qPCR (B) with results shown as mean±sem, n=3, normalised to GAPDH mRNA, or immunoblotting (C) with representative western blot from at least three independent experiments for markers of interferon signaling activation (STAT1 and IRF3 phosphorylation levels), interferon-stimulated genes expression (IRF1 and ISG15), and integrated stress response activation (phosphorylation of eIF2α). Molecular weights are indicated on the right. Band intensities were normalised to that of total protein or GAPDH. Data represents mean ± SEM of three biological replicates. **p<0.01; ***p<0.001; ****p<0.0001; ns: non-significant; using ordinary one-way ANOVA with Šidák’s multiple comparison post-test.

We then quantified the percentage of cells displaying IIR activation following poly(I:C) transfection upon G3Ib or G3Ib’ treatments. Triggering the IIR pathway resulted in IRF3 nuclear translocation upon poly(I:C) stimulation (38.00±4.51%), compared to non-transfected cells (0.67±0.67%). Preventing the formation of G3BP1-foci with G3Ib or the inactive G3Ib’ did does not alter IRF3 nuclear translocation (38.33±5.81% and 36.67±04.10%) (Fig. **3C**). Likewise, IRF1 expression increased upon poly(I:C) stimulus (60.67±3.48%) compared to untreated samples (1.67±0.88%), while no significant changes occurred upon G3Ib or G3Ib’ treatment (66.67±03.23% and 58.00±05.03%) (Fig. **3C**).

These observations are consistent with the unaltered expression of *IFNB1* or ISGs (*ISG56*, *IRF1*, *ISG15*, *IL6*, and *EIF2AK2*) at the mRNA level (Fig. **3D**), or the unchanged levels of IIR-related factors, including IRF3 and STAT1 phosphorylation, as well as ISGs expression (IRF1 and IRF15) (Fig. **3E**). The activation of the ISR, measured by the unaltered levels of eIF2α phosphorylation was also unaffected by G3Ib or G3Ib’ treatment in poly(I:C)-stimulated samples (Fig. **3E**). Overall, these data suggest that the assembly of the poly(I:C)-induced SGs is uncoupled from the induction of intrinsic IIR signalling following viral mimic challenge.

### FMDV infection induces the assembly of G3BP1-foci

Having shown that the assembly of G3BP1-foci does not impact IIR signalling in response to viral mimics, we next addressed the assembly and dynamics of biocondensates formation upon infection of PK-15 cells with FMDV. Previous studies have suggested that FMDV disrupts SG formation, however conflicting data exist regarding the mechanisms involved (25, 26).

To this end, PK-15 cells were infected with FMDV for 3.5 hours and the assembly of G3BP1-foci and FMDV replication monitored using immunofluorescence labelling for G3BP1 and the viral 3A protein, respectively. FMDV infection resulted in the assembly of G3BP1-foci in 52.33±7.31% of the infected cells at 3.5 hours post-infection (hpi) (Fig. **4A-B**), compared to 93.67±0.88% and 27.67±7.69% in cells treated with the positive controls sodium arsenite or poly(I:C), respectively. Interestingly, FMDV-induced G3BP1-foci were present in lower numbers (8.56±2.90) and smaller in size (0.20±0.01 μm^2^) when compared to those induced by sodium arsenite (27.03±1.93, 0.54±0.07 μm^2^) and poly(I:C) (13.87±2.38, 0.33±0.01 μm^2^).

**Figure 4.**
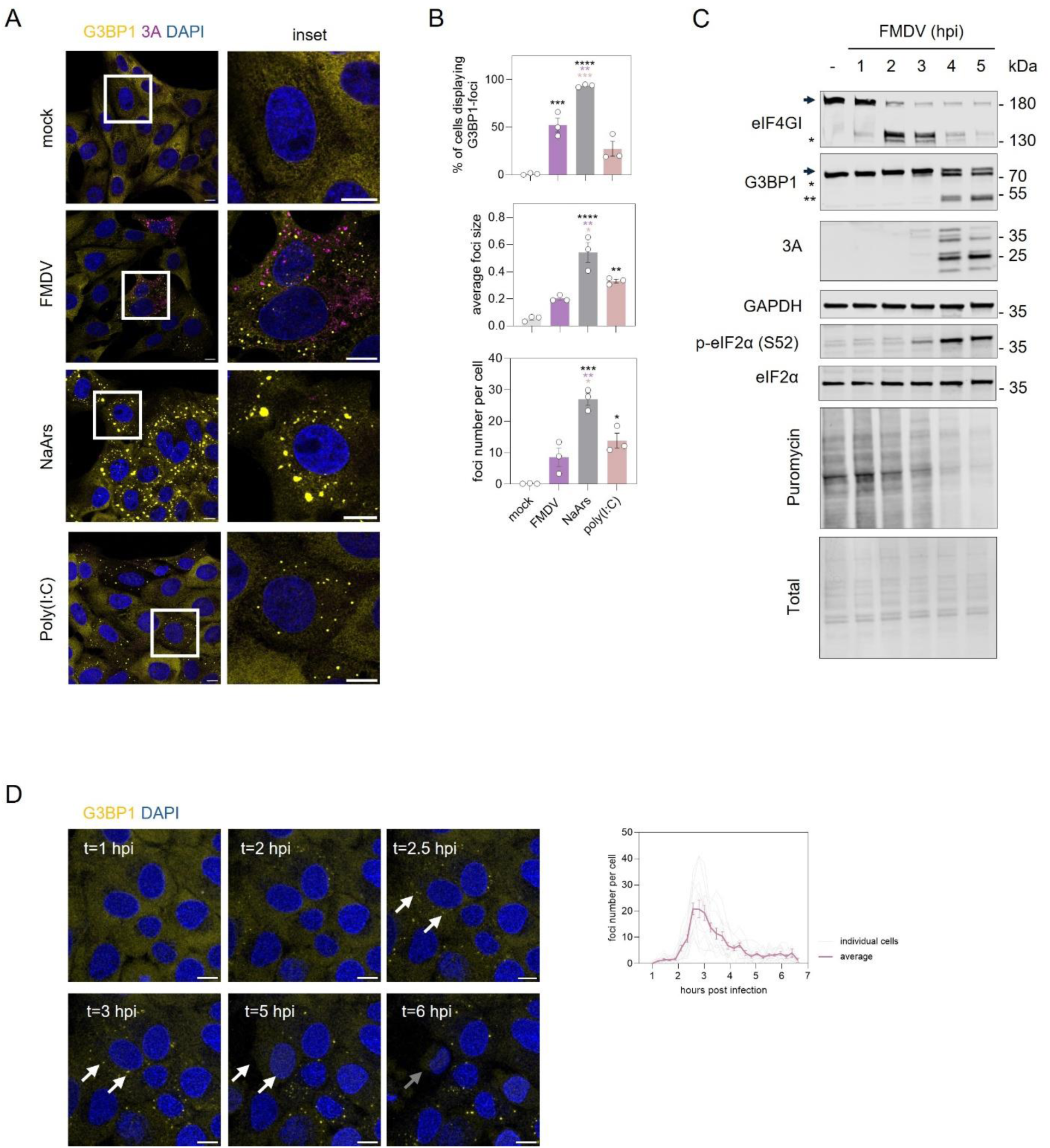
FMDV induces the formation of transient G3BP1-foci during infection. (A-B) PK-15 cells were infected with FMDV for 3.5 hours. Sodium arsenite treatment (1 mM, 1 hour) and poly(I:C) stimulation (1 μg/ml) were used as positive controls. Cells were analysed by immunofluorescence for the SG markers G3BP1 (gold) and FMDV 3A (magenta). Nuclei were stained with DAPI. Scale bars represent 10 μm. The percentage of cells displaying G3BP1-foci was quantified by manual counting of cells of at least 100 cells per replicate. The average size of foci per cell and the number of foci per cell were quantified using the G3BP1 channel and ‘Analyse Particles’ plugin from ImageJ. Results shown as mean ± SEM of at least three independent biological replicates, representing the average of at least 30 cells per condition. *p<0.05; **p<0.01; ***p<0.001; ****p<0.0001; ns: non-significant; using ordinary one-way ANOVA with Šidák’s multiple comparison post-test. (C) Representative western blot from at least three independent experiments for eIF4GI and G3BP1 cleavage, viral infection (FMDV 3A), and integrated stress response activation (phosphorylation of eIF2α and puromycin). Molecular weights are indicated on the right. (D) PK-15 G3BP1:RFP were infected with FMDV and monitored by live-cell imaging for up to 7 hpi. The number of SG per cell of 13 cells that died during this time, reflecting a full cycle of infection, were quantified using the G3BP1 channel and ‘Analyse Particles’ plugin from ImageJ.

FMDV infection results in translational shut-off in infected cells, with evidence of both eIF2α-dependent and independent shut-off signalling (26, 33, 34). To better understand the dynamics of protein synthesis during FMDV infection in PK15 cells, we monitored translation using puromycylation assays and detecting eIF4GI integrity, whose cleavage has been associated with host shut-off during picornavirus infection. FMDV life cycle is rapid with cytophatic effect (CPE) observed after 6 hpi, therefore analyses were carried out hourly up to 5 hpi (Fig. **4C**). Infection was monitored by immunoblotting with an antibody detecting the non-structural viral protein 3A and its polyprotein precursors. Cleavage of eIF4GI was detected as early as 1 hpi, followed by an increase in the eIF2α phosphorylation levels and decrease in translation rates, measured by reduced puromycin intensity, starting at 3 hpi. In addition, G3BP1 cleaved was then detected from 4 hpi.

To follow the kinetics of G3BP1-foci assembly during FMDV infection, we engineered PK-15 cells expressing endogenous RFP-tagged G3BP1, using CRISPR-Cas9-based gene editing. First, we confirmed that these cells respond to FMDV infection similarly to parental PK-15 cells, displaying similar patterns of eIF4GI and G3BP1 cleavage, eIF2α phosphorylation and translational shut off, as well a similar viral replication kinetics (Fig. **S2**). Using this system, we observed the formation of G3BP1-RFP foci, reaching its peak at around 3 hpi (20.77±3.37 foci number per cell) followed by their disassembly and appearance of CPE (Fig. **4D**). These observations further confirm that G3BP1-foci are induced early during FMDV infection and disassemble at later times points, concurrently with G3BP1 cleavage.

### Preventing G3BP1 condensation has no effect on FMDV infection or propagation

To understand the importance of G3BP1-foci assembly for FMDV replication, we treated infected with FMDV at MOI of 1 for 3.5 hours cells in the presence of G3Ib (or G3Ib’ as control) prior to monitoring G3BP1-foci assembly by immunofluorescence. G3Ib treatment resulted in reduced percentage of cells displaying G3BP1-foci from 47.3% to 2.57%, average number per cell from 6.82±2.17 to 0.87±0.57, and average size from 0.19±0.02 to 0.09±0.01 μm^2^, compared to untreated cells, while G3Ib’ had no significant effect (Fig. **5A-B**). Next, we assessed the impact of blocking SG formation on FMDV-induced ISR activation, by measuring the phosphorylation of eIF2α, puromycin levels, and the cleavage of eIF4GI and G3BP1 (Fig. **5C**). G3Ib treatment impaired SG condensation, with no impact on FMDV-induced cleavage of host factors or the activation of ISR when compared to non-treated or G3Ib’-treated samples (Fig. **5C**). To understand the impact of SGs assembly on FMDV replication, PK-15 cells were next infected with FMDV at a low MOI of 0.01 and monitored for signs of CPE, reflected by decrease in cell confluency (35) (Fig. **5D**). Impairing G3BP1-foci assembly did not alter viral replication as the rates of CPE were similar in cells treated with G3Ib,G3Ib’ or untreated cells. The impact on virus replication was investigated by titrating the yield of infection at 6 hpi using plaque assays, revealing similar viral titres and demonstrating no effect of the drugs during infection (Fig. **5E**). Overall, these observations support the hypothesis that FMDV-induced G3BP1-foci do not impact viral replication during infection.

**Figure 5.**
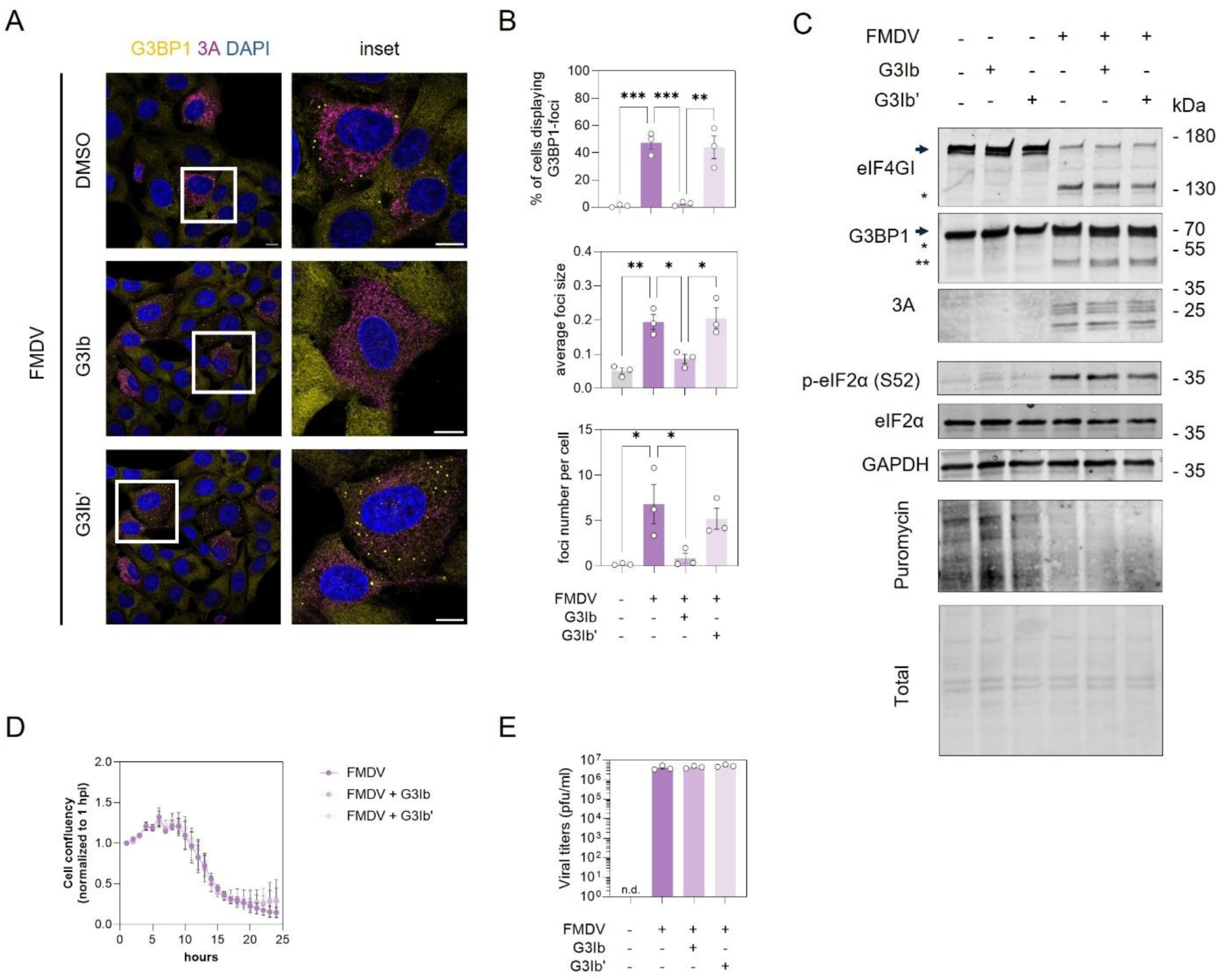
Preventing G3BP1 condensation does not interfere with FMDV replication. (A-C) PK-15 cells were infected with FMDV for 3.5 hours following treatment with G3BP1 condensation inhibitor (G3Ib) or its enactive enantiomer (G3Ib’). (A) Cells were analysed by immunofluorescence for the SG markers G3BP1 (gold) and FMDV 3A (magenta). Nuclei were stained with DAPI. Scale bars represent 10 μm. (B) The percentage of cells displaying G3BP1-foci was quantified by manual counting of at least 100 cells per replicate. The average size of foci per cell and the number of foci per cell were quantified using the G3BP1 channel and ‘Analyse Particles’ plugin from ImageJ. Results shown as mean ± SEM of at least three independent biological replicates, representing the average of at least 30 cells per condition. *p<0.05; **p<0.01; ***p<0.001; ****p<0.0001; ns: non-significant; using ordinary one-way ANOVA with Šidák’s multiple comparison post-test. (C) Representative western blot from at least three independent experiments for eIF4GI and G3BP1 cleavage, viral infection (FMDV 3A), and integrated stress response activation (phosphorylation of eIF2α and puromycin). Molecular weights are indicated on the right. (D) PK-15 cells were infected with FMDV at a MOI of 0.01 in the presence or absence of G3Ib or G3Ib’, alongside a mock-infected controls, and cell confluence was monitored every hour for 24 or 12 hours, respectively, using an IncuCyte Zoom. Data represents an average of three replicates, normalized to the first time-point measured (1 hpi). (E) FMDV titration at 6 hours post-infection following the treatment with G3Ib or G3Ib’. Data represents the average of three independent replicates.

### FMDV-induced G3BP1-foci are not required for IIR activation during infection

Although the assembly of G3BP1-foci does not appear to regulate FMDV replication, this does not exclude a role on IIR modulation during infection. The FMDV leader protease (Lpro) has been previously suggested to counteract SG formation (25) and to contribute to IFN signalling evasion (36–39). To investigate the interplay between ISR and IIR during FMDV infection, we infected PK-15 cells with an engineered FMDV virus lacking the Lpro-encoded region (leaderless FMDV, LL-FMDV) (40). PK-15 cells were infected with WT or LL-FMDV at a MOI of 1 at 3.5 hpi and G3BP1-foci monitored by confocal microscopy. Detailed analysis of foci assembly revealed that LL-FMDV infection resulted in the formation of G3BP1-foci in 67% of infected and 55% of adjacent or bystander cells, whereas the WT virus infection triggered the assembly of G3BP1-foci in 52% of infected but only 8% of bystander cells (Fig. **6A-B**). Analysis of their number and average size revealed that the G3BP1-foci detected in infected cells (i) with WT and LL-FMDV are similar in size (0.21±0.01 μm^2^ and 0.21±0.04 μm^2^) but more abundant in cells infected with LL-FMDV (9.93±0.37 and 24.4±3.8) (Fig. **6C**). Upon LL-FMDV infection, these foci appear as abundant (21.37±3.02) but bigger in bystander cells (b) than in infected cells (i) (0.40±0.06 μm^2^) (Fig. **6C**). This suggests that infection with LL-FMDV induces the assembly of G3BP1-foci with distinct properties in infected and bystander cells.

**Figure 6.**
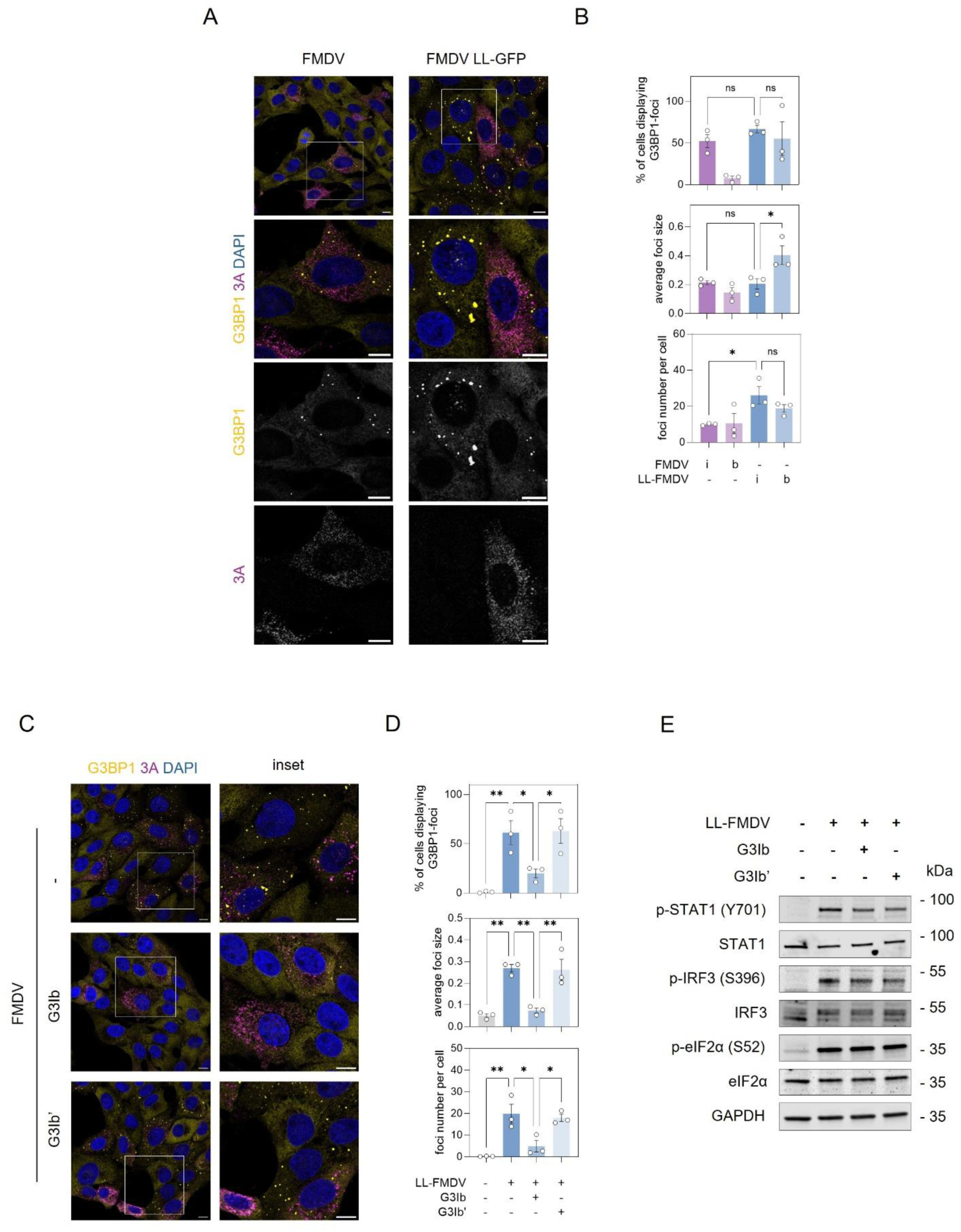
Preventing G3BP1 condensation does not interfere with the activation of interferon response during FMDV infection. (A-B) PK-15 cells were infected with FMDV or LL-FMDV for 4 hours. Cells were analysed by immunofluorescence for the SG markers G3BP1 (gold) and FMDV 3A (magenta). Nuclei were stained with DAPI. Scale bars represent 10 μm. (B) The percentage of cells displaying G3BP1-foci in infected (i) or bystander (b) cells was quantified by manual counting of at least 100 cells per replicate. The average size of foci per cell and the number of foci per cell were quantified using the G3BP1 channel and ‘Analyse Particles’ plugin from ImageJ. Results shown as mean ± SEM of at least three independent biological replicates, representing the average of at least 30 cells per condition. (C-D) PK-15 cells were infected with LL-FMDV for 3.5 hours following treatment with G3BP1 condensation inhibitor (G3Ib) or its enactive enantiomer (G3Ib’). The analysis was done as previously. (E) Representative western blot from at least three independent experiments for innate immune response activation (STAT1 and IRF3 phosphorylation), and integrated stress response activation (phosphorylation of eIF2α). Molecular weights are indicated on the right. *p<0.05; **p<0.01; ***p<0.001; ****p<0.0001; ns: non-significant; using ordinary one-way ANOVA with Šidák’s multiple comparison post-test.

To characterise the dynamics and function of LL-FMDV-induced G3BP1-foci in the activation of IIR, we next treated infected cells with G3Ib or G3Ib’ and analysed both G3BP1-foci formation and IIR activation. First, LL-FMDV-induced G3BP1-foci assembly (61%, 19.88±4.37, 0.27±0.02 μm^2^) was impaired by G3Ib treatment (20%, 4.91±2.67, 0.07±0.01 μm^2^), but not G3Ib’ (63%, 17.98±1.68, 0.26±0.05 μm^2^), in both infected and bystander cells (Fig. **6C-D**). Furthermore, preventing their assembly did not affect either IIR or ISR signalling after 8 hours of infection, with no observed changes in the phosphorylation levels of IRF3 and STAT1, or eIF2α (Fig. **6E**).

Overall, this suggests that the assembly of G3BP1-foci during infection is not important for IIR activation or the IFN-mediated response. In contrast, genetic knock-out of G3BP1/2 impaired both the IFN-mediated response and increased FMDV replication (Fig. **S3**), which suggests a role for G3BP1/2 during infection independently from their condensation properties.

### FMDV-induced G3BP1-foci are *bona fide* SGs

Compositionally heterogenous SGs have been proposed to assemble in response to different stress triggers, with diverse SG-like condensates assembling during viral infection (41). To further characterise FMDV-induced foci, we treated infected cells with cycloheximide, an inhibitor of translation elongation that traps mRNPs into polysomes preventing the formation of canonical SGs (42). Treatment with cycloheximide promoted the disassembly of foci induced by sodium arsenite (from 75.67±3.18% to 6.00±2.08% in non-treated and treated samples, respectively) and infected with FMDV (from 60.33±4.49% to 6.00±1.53%) or LL-FMDV (from 42.67±3.18% to 5.66±0.66%), but not by poly(I:C) stimulation (Fig. **7A-B**). This is supported by a reduction in the number of G3BP1-containing foci following cycloheximide treatment in cells stimulated with sodium arsenite (from 19.167±2.76 to 1.12±0.09) in non-treated and treated samples, respectively) and infected with FMDV (from 8.09±0.59 to 0.53±0.17) or LL-FMDV (from 6.40±1.09 to 0.64±0.), but not by poly(I:C) stimulation (7.65±1.03 and 8.09±0.59). This suggests that G3BP1-foci formed during FMDV and LL-FMDV infection assemble as a consequence of impaired translation, a property of canonical SGs.

**Figure 7.**
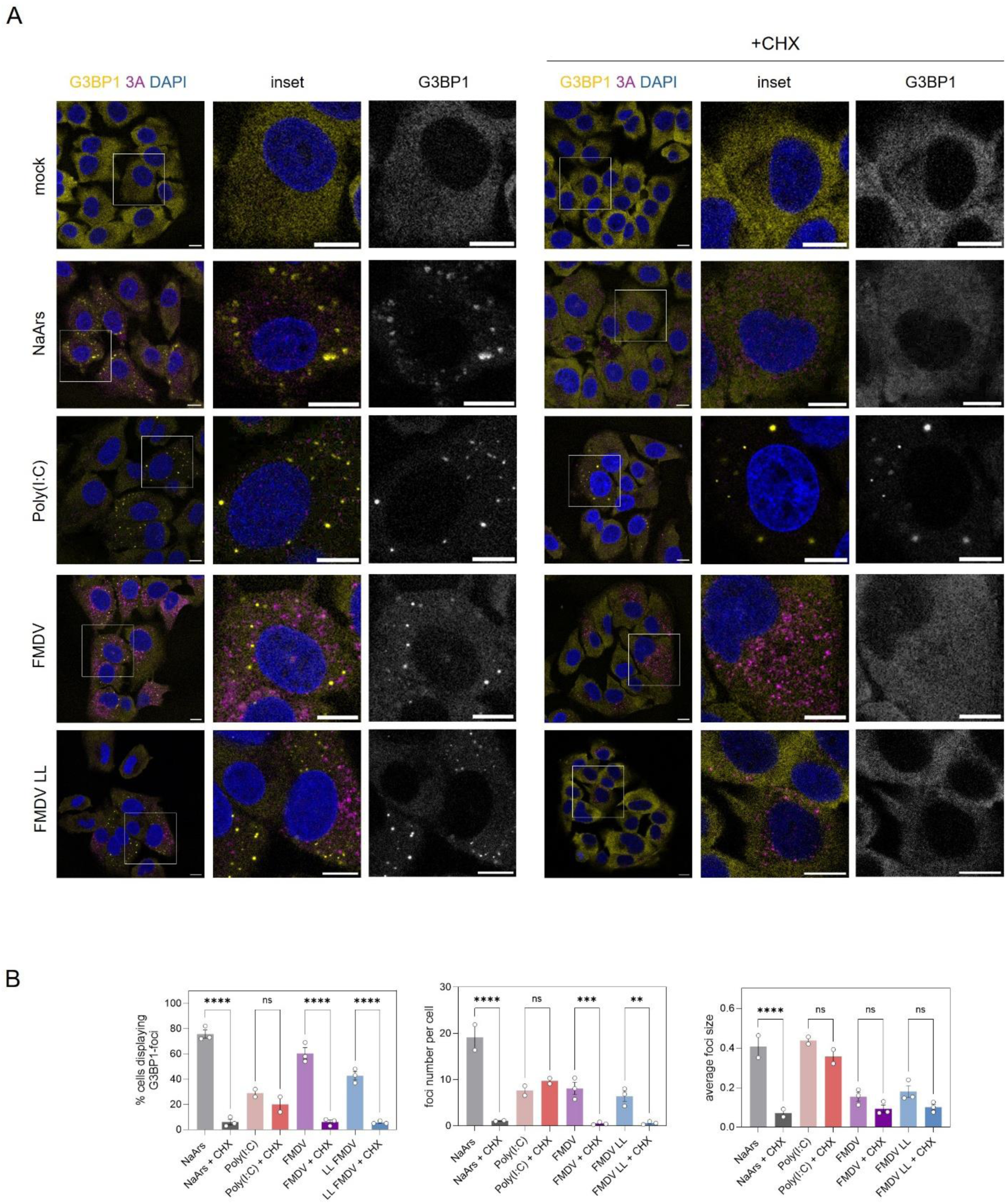
FMDV-induced biocondensates formation are dependent on translation inhibition. PK-15 cells were treated with 1 mM sodium arsenite for 1 h, stimulated with 1 µg/ml poly(I:C), or infected with FMDV or LL-FMDV for 3.5 hours, following by a 30-minute treatment with 25 µg/ml cycloheximide treatment prior to harvesting. (A) Cells were analysed by immunofluorescence for the SG markers G3BP1 (gold) and viral protein 3A (magenta). Nuclei were stained with DAPI. Scale bars represent 10 μm. (B) The percentage of cells displaying G3BP1-foci was quantified by manual counting of at least 100 cells per replicate. The average size of foci per cell and the number of foci per cell were quantified using the G3BP1 channel and ‘Analyse Particles’ plugin from ImageJ. Results shown as mean ± SEM of at least three independent biological replicates, representing the average of at least 30 cells per condition. **p<0.01; ***p<0.001; ****p<0.0001; ns: non-significant; using ordinary one-way ANOVA with Šidák’s multiple comparison post-test.

### Preventing SGs disassembly impairs FMDV replication

Given FMDV infection resulted in early assembly of SGs that later disassemble, we hypothesized SGs disassembly may be important for FMDV replication, and thus that their persistence might impair replication. To test this, we treated PK-15 cells with increasing amount of raloxifene, a drug previously shown to delay SG dissolution (43). In PK-15 cells, concentrations equal or higher to 12.5 μM impaired basal cellular stress state (**Fig. S4A-B**), therefore experiments were performed with doses up to 6.25 μM.

PK-15 were infected with WT FMDV in the presence of raloxifene and SG assembly monitored by immunofluorescence as before (Fig. **8A-B**). Raloxifene treatment increased both the percentage of infected cells displaying SGs from 37.00±2.08% to 66.33±4.84% (Fig. **8B**), and the number of SG per cells, from 3.35±0.35 to 10.26±1.19, while their average size did not change (0.19±0.01 μm^2^ and 0.17±0.01 μm^2^) (Fig. **8B**). CPE was measured after infecting PK-15 cells with FMDV at a MOI of 0.01 in the presence of raloxifene (Fig. **8C**). Full CPE effect was observed for FMDV-infected cells, untreated or treated with the lowest concentration of raloxifene tested (1.56 μM) at around 14 hpi, whereas treating cells with 3.12 or 6.25 μM at around 16 and 18 hpi, respectively, resulted in reduced CPE indicating attenuated viral replication.

**Figure 8.**
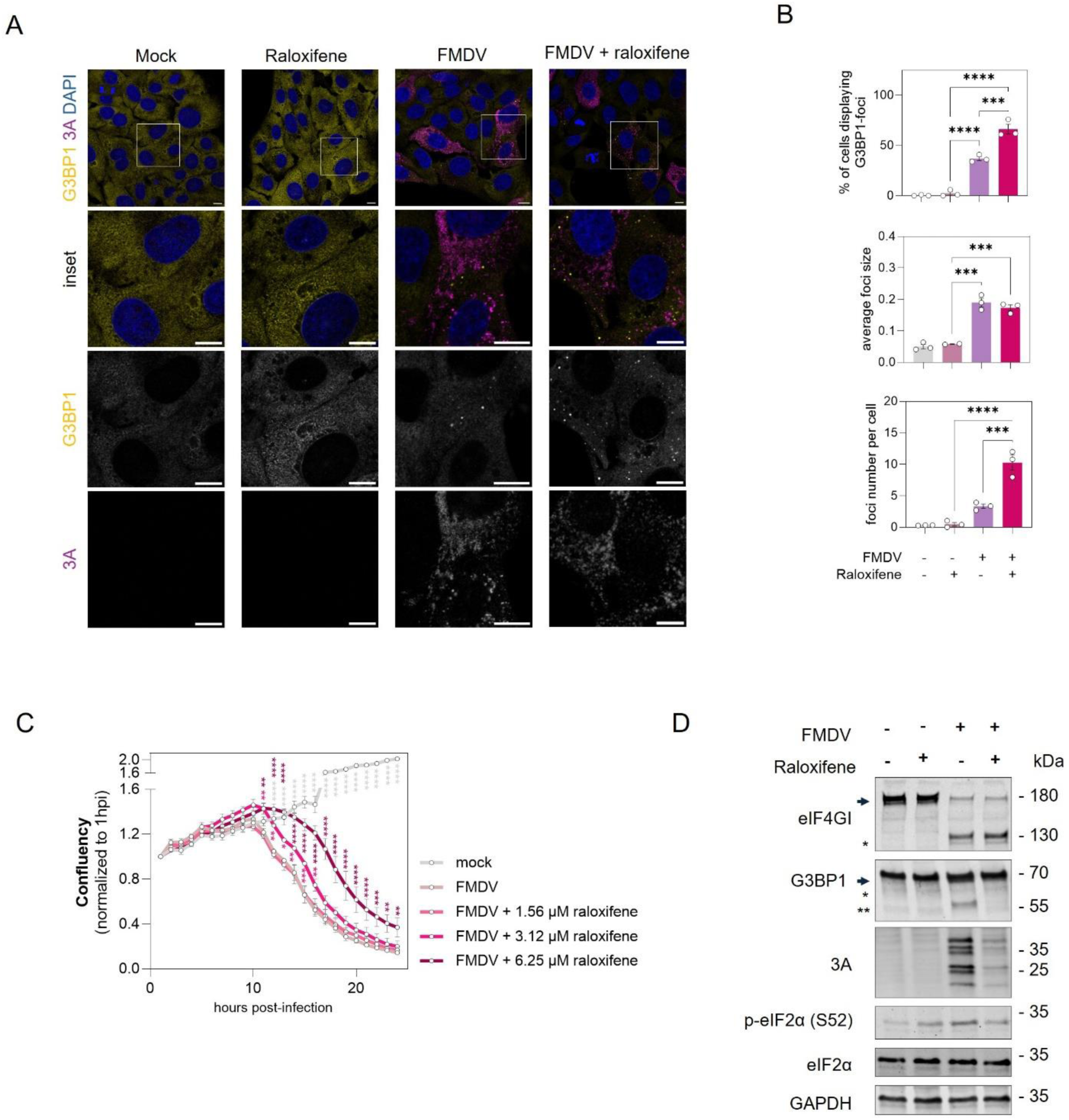
Preventing SGs disassembly halts viral replication. (A-C) PK-15 cells were infected with FMDV for 4 hours with or without 6.25 µM raloxifene. (A) Cells were analysed by immunofluorescence for the SG markers G3BP1 (gold) and viral protein 3A (magenta). Nuclei were stained with DAPI. Scale bars represent 10 μm. (B) The percentage of cells displaying SGs was quantified by manual counting of cells with G3BP1-foci of at least 100 cells per replicate. The average size of foci per cell and the number of foci per cell were quantified using the G3BP1 channel and ‘Analyse Particles’ plugin from ImageJ. Results shown as mean ± SEM of at least three independent biological replicates, representing the average of at least 30 cells per condition. ***p<0.001; ****p<0.0001; ns: non-significant; using ordinary one-way ANOVA with Šidák’s multiple comparison post-test. (C) PK-15 cells were infected with FMDV at a MOI of 0.01 with increasing levels of raloxifene, alongside a mock-infected control, and cell confluence was monitored every hour for 24 hours using an IncuCyte Zoom. Data represents an average of three replicates, normalized to the first time-point measured (1 hpi). *p<0.05, **p<0.1, *** p<0.001, ****p<0.0001; using two-way ANOVA with Šidák’s multiple comparison between mock-infected or FMDV-infected cells treated with raloxifene and FMDV-infected cells. (D) Representative western blot from at least three independent experiments for eIF4GI and G3BP1 cleavage, viral infection (FMDV 3A), and integrated stress response activation (phosphorylation of eIF2α). Molecular weights are indicated on the right.

Next, we analysed the effect of raloxifene on FMDV replication and cleavage of eIF4GI and G3BP1. PK-15 cells were infected with FMDV at a MOI of 1 for 3.5 hours and either untreated or treated with 6.25 μM raloxifene at the time of infection. Raloxifene treatment reduced the amount of viral products (Fig. **8D**), suggesting impaired viral replication. The cleavage of eIF4GI, previously described as Lpro-driven (44, 45), was unaltered by raloxifene (Fig. **8D**), suggesting no effect on Lpro activity, while in contrast, G3BP1 cleavage was reduced (Fig. **8D**), supporting a role for 3Cpro, rather than Lpro, in G3BP1 proteolytic targeting during infection.

Overall, this suggests that inducing persistent condensates does not impact ISR signalling or host shut-off, but alters FMDV replication kinetics, suggesting the disassembly of G3BP1-foci is a process that can be targeted to attenuate FMDV infection.

## Discussion

Biocondensates formation has emerged as a key process triggered in response to various physiological and pathological challenges to help cells rapidly adjust fundamental processes (46). Distinct biocondensates with variable functions, including SGs, paracrine granules, RLBs, dsRNA-induced foci (dRIFs), or viral aggregated RNA condensates (VARCs), have been shown to assemble in response to viral infections, with conflicting literature on their roles during infection (41). Understanding the composition and dynamics of biocondensates is essential to uncover their biological function and relevance for antiviral responses or viral replication.

SGs are among the most studied condensates and have been suggested to act as platforms for innate immune activation in response to viral infection, providing a scaffold for nucleic acids recognition, or activation of antiviral signalling (16, 23, 47, 48). However, recent studies challenged this by demonstrating that SGs do not concentrate antiviral proteins, nor regulate the activation of cell intrinsic antiviral signalling pathways (24, 49). Here, we used distinct viral RNA mimics, and FMDV to establish that SGs assembly is uncoupled from the activation of IIR signalling in porcine cells. Our work demonstrates that triggering RIG-I/MAVS-dependent interferon signalling with the RIG-I ligand 3p-hpRNA is not sufficient to promote SG formation. We also show that the formation of condensates following poly(I:C) stimulation occurs concomitantly to the activation of IRF3-mediated antiviral signalling and that it further regulates the synthesis of ISGs, such as IRF1. Cells that assemble condensates show reduced levels of IRF1, likely due to a halted translation state despite active mRNA transcription. A previous report has suggested that the *IFNB1* mRNA is stalled within SGs of translationally arrested cells (50), which suggest other ISG mRNAs might also undergo a similar process.

To clarify the importance of these condensates for IIR activation, we show for the first time that chemically impairing the assembly of G3BP1-foci with the condensation inhibitor G3Ib (32) does not impact IIR activation. By preventing G3BP1 self-assembly, G3Ib disrupts the further recruitment and accumulation of translation initiation factors or other SG proteins (e.g., eIF3η, PABPC1) and ultimately the assembly of G3BP1-foci, without affecting its RNA-binging activity or the cellular translation initiation rates (32). This indicates that, in our model, physically preventing the condensation of G3BP1-foci does not impact IIR signalling.

To understand the interplay between G3BP1-foci and IIR activation during viral infection, we focussed our studies on FMDV, a virus previously shown to exploit SGs assembly and evade RLR-mediated IIR (25, 36, 37, 39). We characterized the dynamics of biocondensates assembly and disassembly in porcine epithelial cells during infection. Live-cell imaging revealed that G3BP1-foci form early during infection, peaking roughly at 3 hpi, before disassembling prior to initiation of CPE at around 6 hpi. This assembly peak also corresponds to the initial timing of eIF2α phosphorylation, which is followed by a later cleavage of G3BP1, suggesting that this might be the trigger for their disassembly. Our data reconciliate previous reports showing that FMDV modulates SGs formation via G3BP1 cleavage (25, 26) and further establish that FMDV promotes SGs disassembly rather than preventing their formation by monitoring their kinetics during infection. These observations are supported by the morphological characterization of the FMDV-induced SGs, which are smaller than the canonical SGs induced by sodium arsenite and resemble the round small foci induced by poly(I:C). Yet, poly(I:C) and FMDV-induced SGs have distinct features and potentially biological roles, given of differences in sensitivity to cycloheximide, suggesting that the latter share some attributes with canonical SGs.

To study the relevance of FMDV-induced SGs during infection on the IIR activation, we took advantage of a FMDV mutant lacking the leader protein (LL-FMDV), which has been shown to downregulate the RIG-I/MDA5 pathway and interfere with the activation of downstream antiviral effectors to impair the IFN-mediated immune response (36, 37, 40). FMDV Lpro has also been suggested to target G3BP1/2 to suppress SGs (25). Infection with LL-FMDV allowed us to probe the importance of biocondensates, while the IIR is active. We observed the formation of granules displaying distinct morphology in infected or bystander cells, which paves the way for thorough compositional characterization, to unravel putative differences in biogenesis and/or biological functions. Interestingly, we previously proposed the transfer of stress signalling to bystander cells as a cellular mechanism to limit viral propagation (13). One can speculate that Lpro might be involved in suppressing a paracrine mechanism that triggers an antiviral state in bystander cells, leading to the formation of biocondensates.

We further show that chemically preventing G3BP1 condensation during infection does not modulate FMDV replication, neither modulates the IIR in LL-FMDV-infected cells, supporting our previous observations that SGs formation and IIR are uncoupled. Nevertheless, FMDV remains able to cleave G3BP1 when SG assembly is prevented, which together with increased replication in G3BP1/2 KO cells suggests that FMDV may have evolved to counteract G3BP1 functions that are independent from SG formation. These are all compelling arguments supporting that G3BP1 exert antiviral functions, independently from its condensation properties. This is supported by previous reports of G3BP1 involvement in promoting the IIR to viral DNA or RNA, and that the absence of G3BP1 in mammalian cells alters IIR activation levels following a dsRNA stimulus (23, 51, 52).

Other members of the *Picornaviridae* family have been shown to induce SG formation and cleave G3BP1. Seneca Valley virus (SVV), which also causes vesicular disease in pigs, transiently induces SG formation at earlier stages of infection, in a PKR-eIF2α dependent manner, and inhibits SGs at later stages through 3C-mediated disruption of eIF4GI-G3BP1 interaction (53). However, genetically preventing SGs formation did not affect SVV propagation (53). Similar dynamics are observed in poliovirus (54), coxsackievirus B3 (CVB3) (55), and enterovirus 71 (56, 57), which all induce SGs at an early stage but limit SGs as infection progresses, a trait associated with 3Cpro-mediated G3BP1 cleavage.

Lastly, to explore whether preventing G3BP1-foci disassembly would interfere with viral infection, we treated the infected cells with raloxifene, a drug previously shown to delay the disassembly of hypoxia-induced SGs (43). Our data show that raloxifene can delay the disassembly of FMDV-induced G3BP1-foci resulting in delayed infection, making this process a potential target to block FMDV replication. Interestingly, raloxifene was previously shown to exert an antiviral effect against SARS-CoV-2 (58), influenza A virus (59), hepatitis C virus (60); however, the underpinning cellular mechanism is yet poorly understood. With previous therapeutic approaches identifying drugs hardening RSV-induced condensates to impair replication, this opens the door for pharmacological targeting of SGs during FMDV infection (61).

In conclusion, we established for the first time in a porcine cell model that the antiviral response to dsRNA or an RNA virus infection and the assembly of biocondensates – including canonical FMDV-induced SGs or cycloheximide-resistant poly(I:C)-driven condensates – are triggered independently, and we further suggest the targeting ofG3BP1-foci disassembly as a potential antiviral target against FMDV.

## Materials and Methods

### Cell culture

Information about the cell lines used (PK-15 and BHK-21) and engineered (PK-15 G3BP1-RFP and PK-15 G3BP1/2 KO) is provided in **Text S1** in the supplemental material.

### Chemical treatments

Detailed information about the chemicals used in the experiments, including sodium arsenite, poly(I:C), 3p-hpRNA, G3Ib, cycloheximide, and raloxifene, can be found in **Text S1**.

### FMDV production, infection and titration

Infections were performed with a cell line-adapted strain of FMDV (O1K) and a mutated GFP-tagged leaderless FMDV (LL-FMDV). Detailed protocols regarding production of virus stocks, titration, infection, and CPE development, is provided in **Text S1**.

### Immunofluorescence microscopy and live cell experiments

For immunofluorescence assays, PK-15 or PK-15 G3BP1/2 KO cells were seeded on sterilized coverslips and fixated with paraformaldehyde after experiments, whereas for live experiments, PK-15 G3BP1-RFP cells were seeded into chambered cover glasses and imaged for up to 6 hours after infection. Thorough information regarding protocol, antibodies, and samples imaging and analysis, can be found in **Text S1**.

### Immunoblotting and RT-qPCR

Cells were seeded in 24-well plates to be around 80% confluent at the indicated times. Detailed information regarding immunoblotting and RT-qPCR, including samples processing, the methods used, antibodies, and primers, are provided in **Text S1**.

### Ribopuromycyclation assay

To capture translation efficacy, cells were treated with 10 μg/ml of puromycin (Sigma-Aldrich) for 5 min at 37 C to label the nascent polypeptide chains before addition of 180 μM of emetine (Sigma-Aldrich) to block the translation elongation with a further incubation of 2 min at room temperature. Puromycin incorporation was quantified by immunoblotting or immunofluorescence as described before (13).

### Statistical analyses

Statistical analyses were performed using GraphPad Prism software (Ver 10) and at least three biological replicates were analysed in the experiments. Statistical tests performed and correspondent significance are indicated in the figure legends.

## Acknowledgments

Work in N.L.’s laboratory is supported by the Biotechnology and Biological Sciences Research Council institute strategic program grant (BBS/E/PI/230002), national bioscience research infrastructures for high and low containment services and science platforms (BBS/E/PI/23NB0004 and BBS/E/PI/23NB0003), and research grant (BB/X018431/1). Work in A.R.’s laboratory was supported by the Deutsche Forschungsgemeinschaft (DFG, German Research Foundation) project number 240245660 SFB1129 TP13.

## Data availability

All relevant data can be found within the article and its supplementary information.

## Supplemental Figures

**Figure S1.**
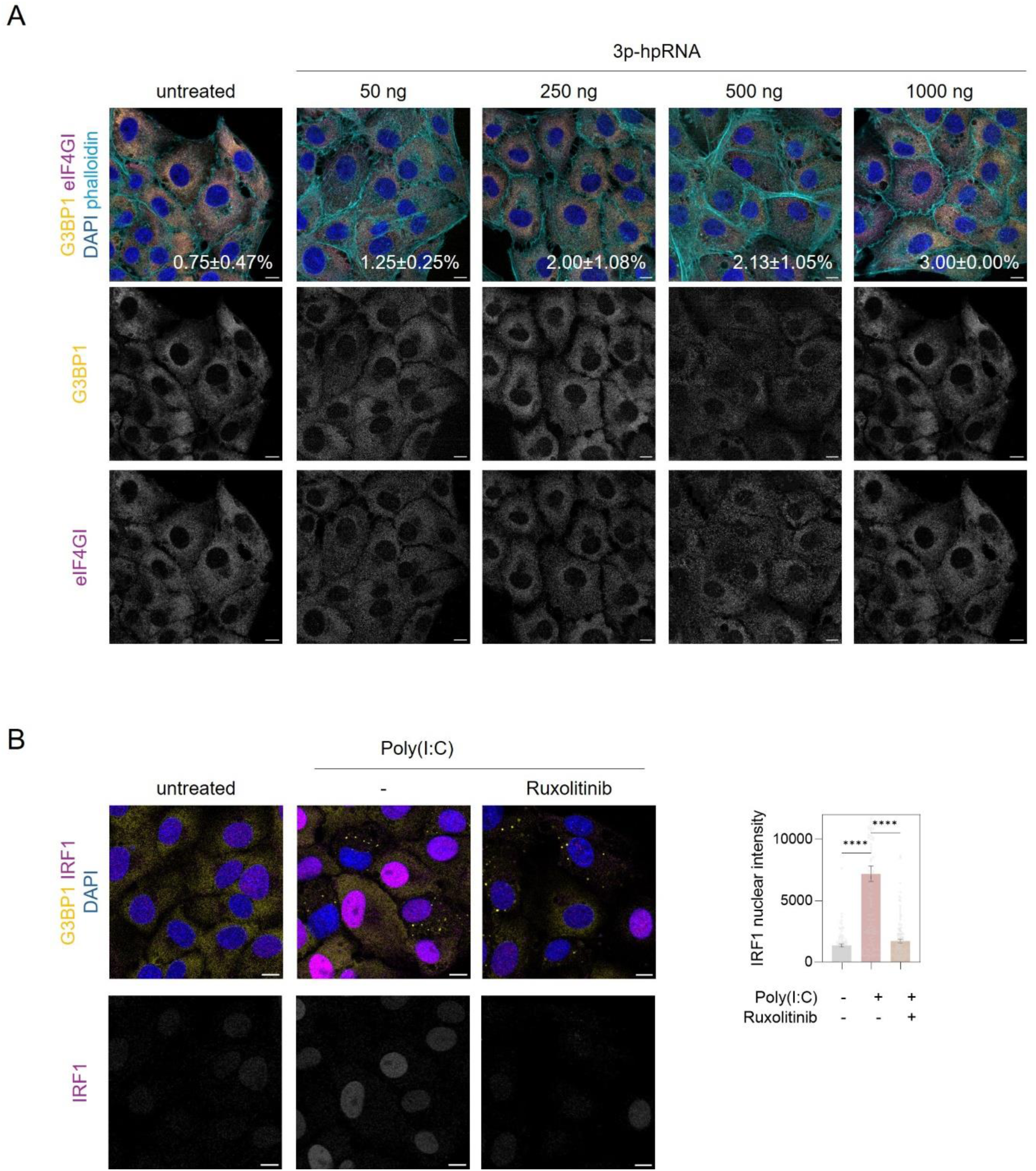
RIG-I/MAVS activation does not induce G3BP1-foci formation. (A) PK-15 cells were stimulated with increasing concentrations of 3p-hpRNA for 6 h. Cells were analysed by immunofluorescence for the SG markers G3BP1 (gold) and eIF4GI (magenta), and F-actin marker phalloidin (cyan). Nuclei were stained with DAPI. Scale bars represent 10 μm. The percentage of cells displaying SGs was quantified by manual counting of at least 100 cells per replicate. **IRF1 expression depends on interferon-mediated signalling.** (B) PK-15 were stimulated with poly(I:C) for 6 h in the presence of ruxolitinib, an inhibitor if interferon signalling. Cells were analysed by immunofluorescence for the SG marker G3BP1 (gold) and innate immune response activation marker IRF1 (magenta). Nuclei were stained with DAPI. Scale bars represent 10 μm. Quantification of IRF1 nuclear intensities was done using ImageJ considering at least 100 cells per sample. ****p<0.0001; using ordinary one-way ANOVA with Šidák’s multiple comparison post-test.

**Figure S2.**
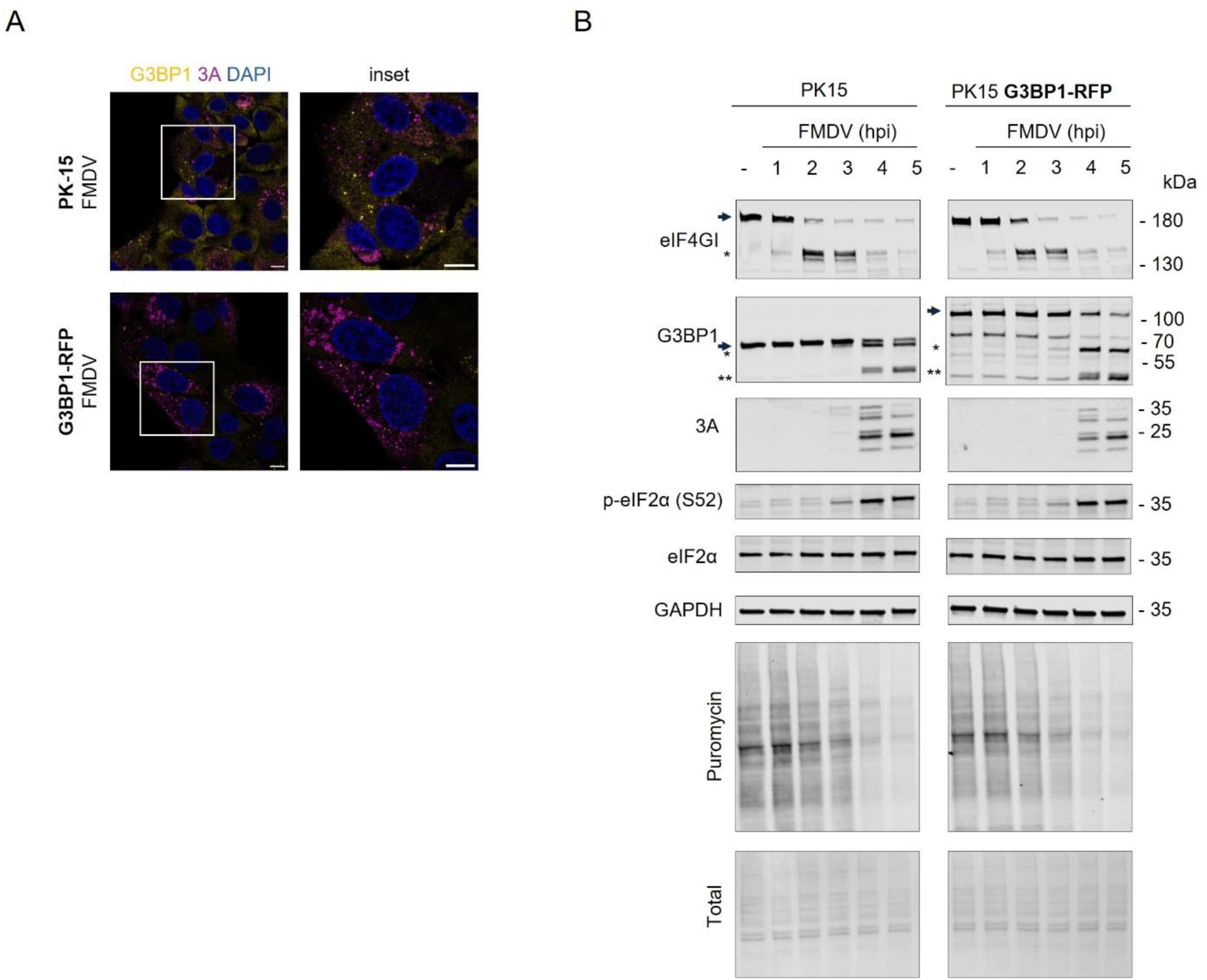
Comparison of infection-mediated cell signalling between PK-15 and PK- 15 G3BP1:RFP. (A) PK-15 and PK-15 G3BP1:RFP cells were infected with FMDV for 3.5 hours. Cells were analysed by immunofluorescence for the SG markers G3BP1 (gold) and FMDV 3A (magenta). Nuclei were stained with DAPI. Scale bars represent 10 μm. (B) Representative western blot from at least three independent experiments for eIF4GI and G3BP1 cleavage, viral infection (FMDV 3A), and integrated stress response activation (phosphorylation of eIF2α and puromycin). Molecular weights are indicated on the right.

**Figure S3.**
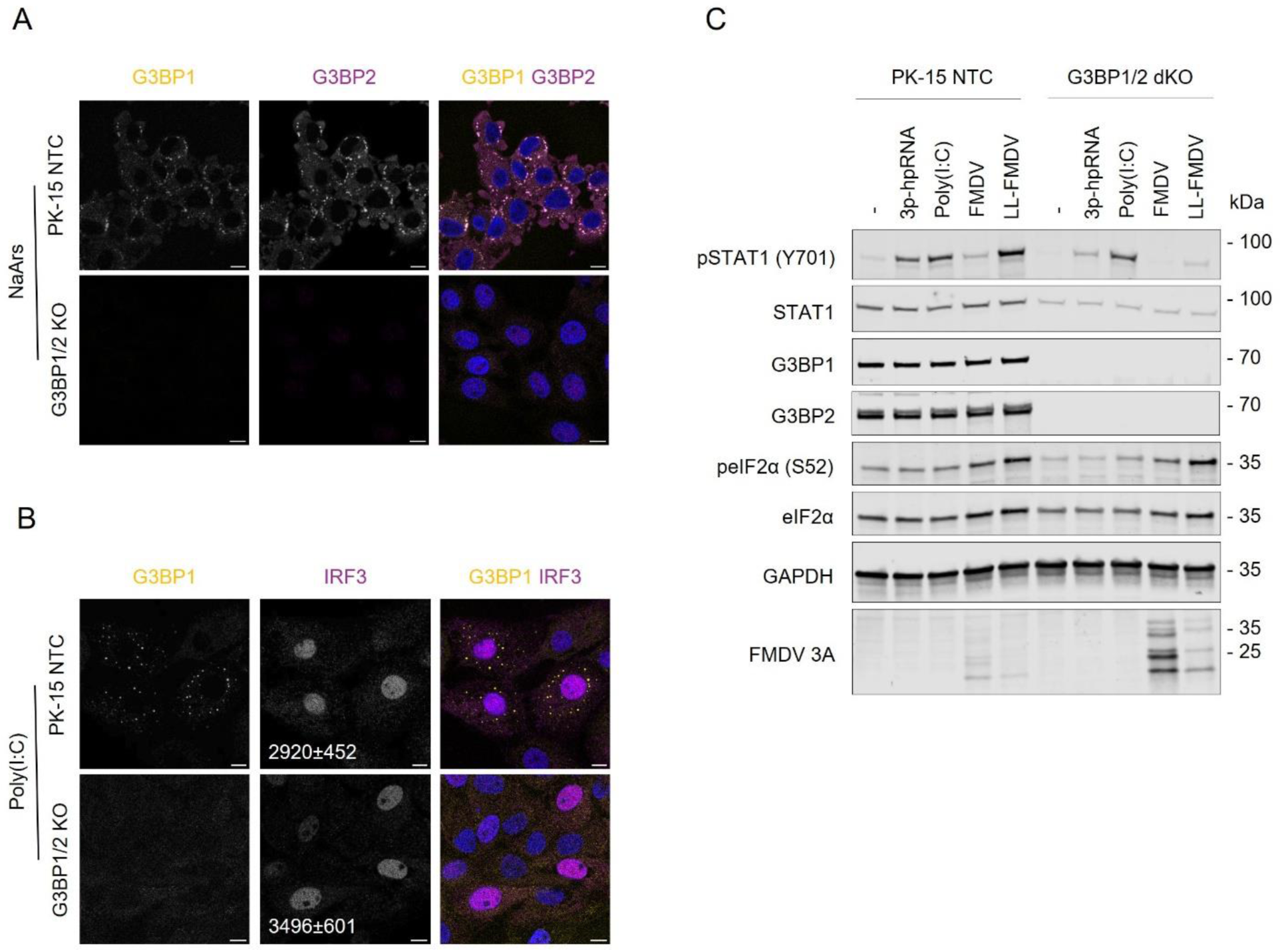
FMDV replicates better in the absence of G3BP1/2 due to a limited interferon-mediated response. (A-B) PK-15 NTC or G3BP1/2 KO cells were (A) treated with 1 mM sodium arsenite for 1 h or (B) stimulated with 1 µg/ml poly(I:C) for 4 h. Cells were analysed by immunofluorescence for the SG markers G3BP1 (gold) and (A) G3BP2 (magenta) or (B) innate immune response activation markers IRF3 (magenta). Nuclei were stained with DAPI. Scale bars represent 10 μm. (C) PK-15 NTC or G3BP1/2 KO cells were stimulated with 1 µg/ml 3p-hpRNA or poly(I:C), or infected with FMDV or LL- FMDV for 4 hours. Western blot for innate immune response (STAT1 phosphorylation) and integrated stress response activation (phosphorylation of eIF2α), G3BP1/2 levels, and FMDV 3A. Molecular weights are indicated on the right.

**Figure S4.**
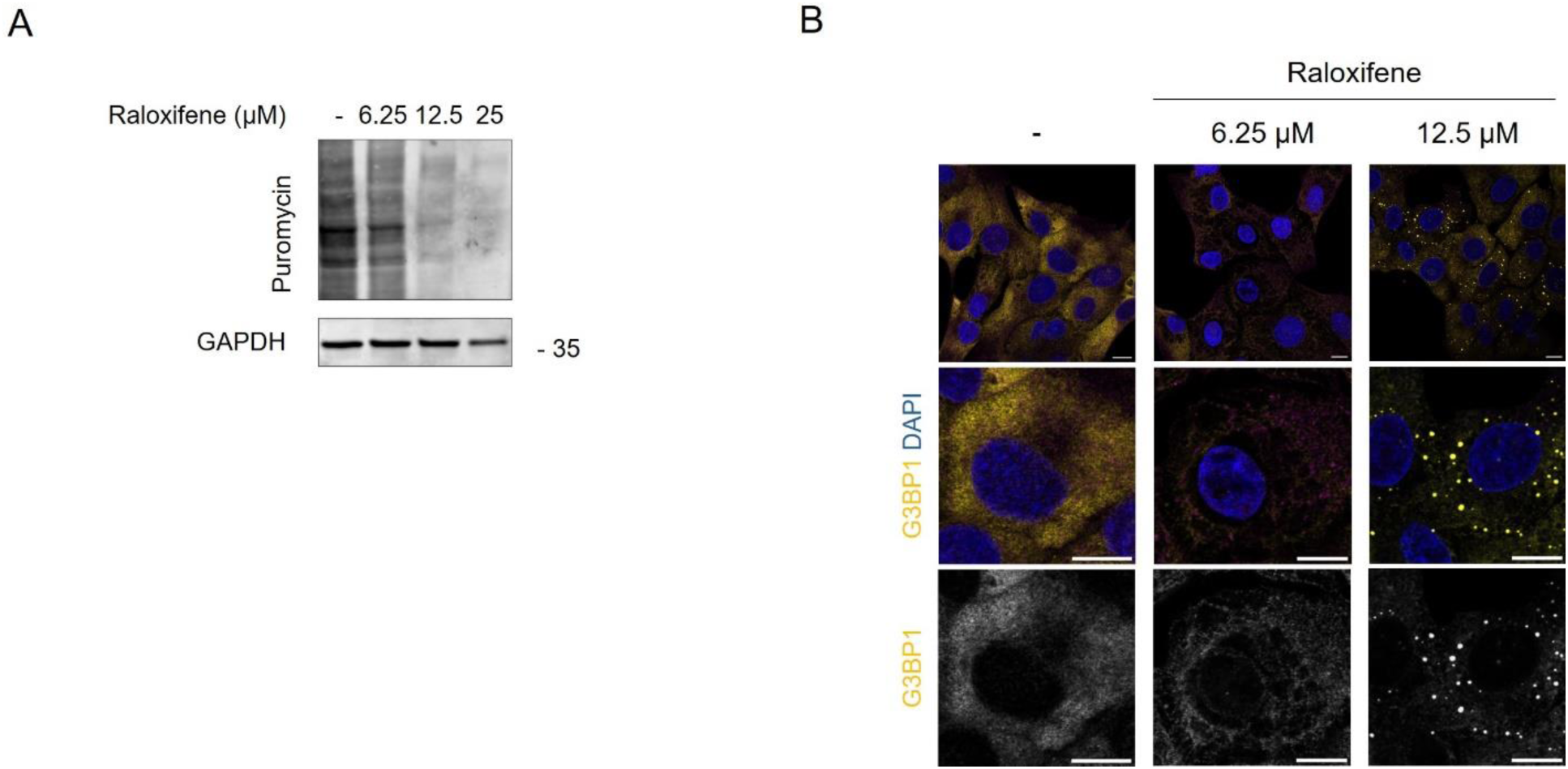
Concentrations of raloxifene higher than 6.25 µM induce translational shut-off and stress granules formation. (A-B) PK-15 cells were treated with increasing concentrations of raloxifene for 4 hours. (A) Representative western blot from at least three independent experiments for puromycin, indicating translation levels. Molecular weights are indicated on the right. (B) Cells were analysed by immunofluorescence for the SG marker G3BP1 (gold). Nuclei were stained with DAPI. Scale bars represent 10 μm.

## Text S1 – Supplemental Material and Methods

### Cell culture

PK-15 cell lines were grown in Dulbecco’s modified Eagle’s medium (DMEM, Sigma-Aldrich, #D5796) supplemented with 10% foetal bovine serum (FBS) (Life Science Production), 100 U/ml penicillin and 100 μg/ml streptomycin (Sigma Aldrich), and 1 mM sodium pyruvate (Gibco), in a humidified incubator at 37°C and 5% CO2 environment. BHK-21 cells were grown in Glasgow’s MEM (GMEM, Sigma-Aldrich, #G5154) supplemented with 10% FBS, 100 U/ml penicillin and 100 μg/ml streptomycin, 2mM L-glutamine (Sigma-Aldrich), and 5% Tryptose Phosphate Broth (TPB, Sigma Aldrich, #18050-039).

PK-15 G3BP1-RFP cells were generated following the guidelines of the TrueTag™ Donor DNA Kit, RFP using a custom gRNA (5’-CGAAGAGCAATTCACTGCCT-3’) and homology arm primers, forward 5’-CAGGATTTGGAGTGGGAAGGGGGCTTGCGCCACGGCAGGGAAGTGGCTCAGGTT CTGGA-3’ and reverse 5’-GACAGGGTTTGTATTGTTGCGCGAAGAGCAATTCACTTGGCCGATCGCATACAGAG-3’, targeting the C-terminal of porcine G3BP1 (*Sus scrofa*, NM_001205405.1). Cells were grown with complete DMEM supplemented with 5 μg/ml of blasticidin (InvivoGen) to ensure selection. Single cell clones were expanded and selected for RFP and G3BP1 assembly following stress treatments. To proceed with the live-cell experiments, four clones were pooled together in equal proportions.

PK-15 G3BP1/2 KO cells were generated using a three-gRNA strategy to delete a portion of each locus encoding the corresponding protein. gRNAs (Table **1**) (Synthego) were resuspended in 1× TE buffer (10 mM Tris-HCl 0.1 mM EDTA, pH 7.5) to a final concentration of 100 μM (100 pmol/μl). The assembly of gRNA/Cas9 ribonucleoprotein complexes (RNPs) were obtained by gently mixing gRNAs with recombinant Alt-R Cas9 protein (Integrated DNA Technologies, IDT) at a 9:1 ratio in Nucleofector solution from the SF Cell Line Nucleofector Kit (Lonza), followed by 10 min incubation at room temperature. PK-15 cells (1E+06) were resuspended in 25 μl of Nucleofection solution and mixed with 25 μl of each pre-assembled RNPs. Nucleofection was performed using an Amaxa 4D nucleofector (Lonza). Nucleofected cells were incubated for 48 h prior to clonal amplification and screening for homozygous clones using target-specific PCR (GoTaq Hot Start, Promega) on genomic DNA using the correspondent primers (Table **2**). After genotyping, expression levels of each protein in controls and KO single-cell clones were assessed by western blotting. After characterization and for further experiments, cell pools were generated by mixing three clones in equal proportions.

**Table 1.**
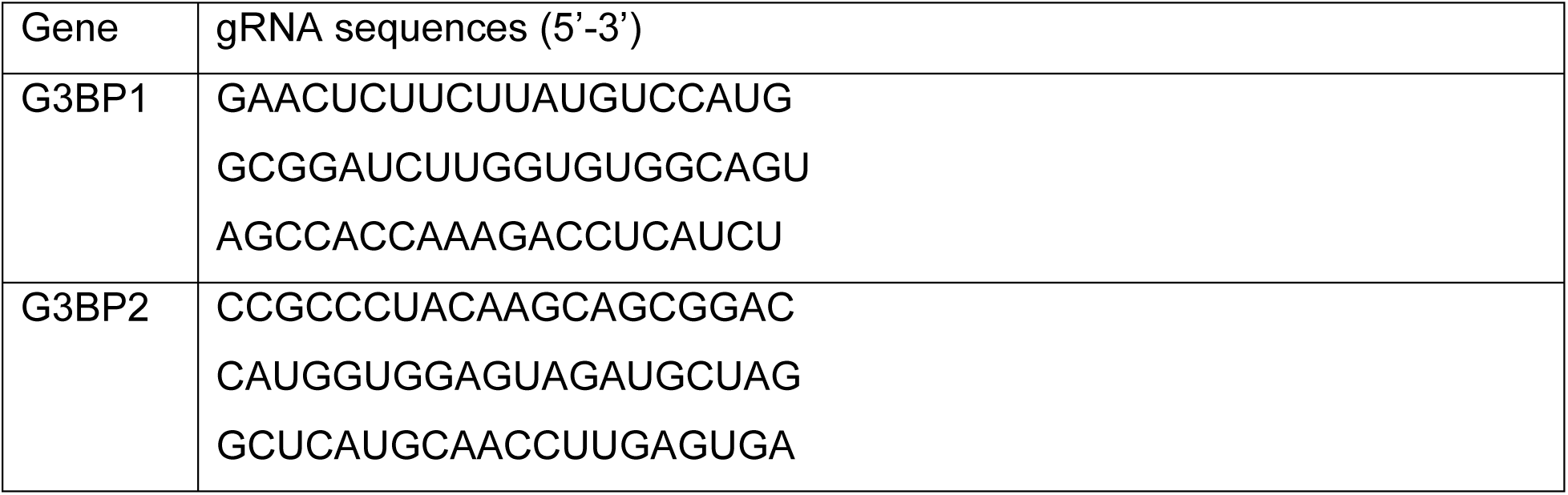
gRNA sequences used for the KO cells.

**Table 2.**
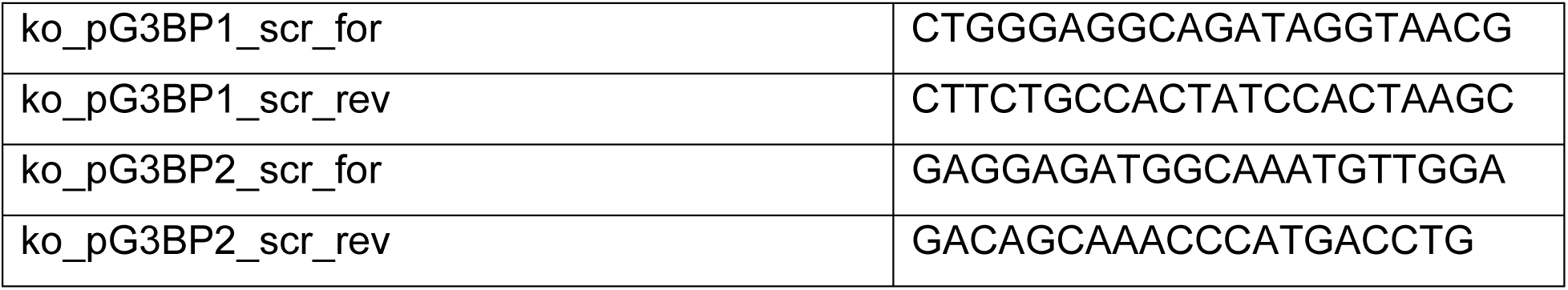
Primers used for the characterization of KO cells.

### Chemical treatments

Sodium arsenite (Sigma-Aldrich) was diluted in H2O and used at a concentration of 1 mM for 1 h. Poly(I:C) (Sigma-Aldrich) diluted in H2O, 150 mM NaCl, and 3p-hpRNA (InvivoGen) diluted in nuclease-free water, were transfected using lipofectamine 2000 (Thermo Fisher Scientific) at 1:3 in Opti-MEM (Thermo Fisher Scientific), according to manufacturer’s instructions. G3Ib or the inactive form G3Ib’, were reconstituted in DMSO and added to the cells 20 minutes prior to treatments or at the time of the infection, at a concentration of 50 µM. Cycloheximide (Sigma Aldrich) was reconstituted in ethanol and added to the cells at 25 μg/ml, 30 minutes prior to sample harvesting. Raloxifene (Cambridge Bioscience) was reconstituted in DMSO and added to the cells at the time of the infection at 1.5 – 6.25 µM.

### FMDV infections

FMDV and LL-FMDV viruses were recovered from BHK-21 cell after full CPE in GMEM supplemented with 1% FBS (Thermo Fisher), 100 U/ml penicillin and 100 μg/ml streptomycin, 2 mM L-glutamine, and 5% TBP. Viral titres were then determined by plaque assays. BHK-21 cell monolayers were infected with 10-fold serial dilutions of virus stock, overlaid with Eagle overlay media supplemented with 5% TBP 100 U/ml penicillin and 100 μg/ml streptomycin (Sigma Aldrich), and 0.6% Indubiose (MP Biomedicals), and incubated for 72 hours at 37°C. Cells were fixed and stained with 1% (w/v) methylene blue in 10% (v/v) ethanol and 4% formaldehyde in PBS. PK-15 cells were seeded in 24-well plates and inoculated with 150 μl of virus dilution in DMEM supplemented with 1% FBS, 100 U/ml penicillin and 100 μg/ml streptomycin, 2 mM L-glutamine and 1 mM sodium pyruvate. After 30 minutes, the inoculum was removed and 500 μl of fresh media were added to the cells with the correspondent treatment, when mentioned. The development of CPE was monitored every hour using the Incucyte S3 Live-Cell Analysis System (Essen BioScience) and imaged using phase light every four hours until complete CPE was reached (35).

### Immunofluorescence microscopy and live cell experiments

Cells were seeded on sterilized coverslips in a 24-well plate and treated as required. Media was removed and fixed with 4% paraformaldehyde (PFA) in PBS for 20 min at RT. Fixation solution was removed, coverslips were rinsed with PB and stored at 4 °C or processed immediately. Cells were permeabilized with 0.1% Triton X-100 (Sigma-Aldrich) in PBS for 5 min at RT and blocked with 0.5% bovine serum albumin (Sigma-Aldrich) in PBS for 1h at RT. Cells were then incubated with primary antibody (Table **3**) for 1h at RT, washed 3 times with PBS and incubated with secondary antibody solution containing 0.2 μg/ml DAPI for 1h at RT. Secondary antibodies used were goat anti-rabbit or goat anti-mouse Alexa 488, 555 or 647 (1:500, Invitrogen). When mentioned, phalloidin 647-conjugated (1:500, SC-363797) was also added. Coverslips were then rinsed 3 times with PBS and mounted onto microscope slides with 5 μl Mowiol 4-88 (Sigma-Aldrich). Cells were visualized and imaged with Leica STELLARIS 5 inverted microscope or Leica SP8 CLSM upright microscope using a 60X objective. Confocal images were analysed utilizing Image J.

**Table 3.** Antibodies used on immunofluorescence staining.

| Primary antibodies used in WB | Host species | Reference |
| --- | --- | --- |
| IRF3 | Rabbit | Cell Signalling Technology, #8478 |
| IRF1 | Rabbit | Cell Signalling Technology, #8784 |
| Puromycin | Mouse | Millipore, MABE343 |
| G3BP1 | Rabbit | Merck, G6046 |
| G3BP1 | Mouse | BD Transduction, #611127 |
| FMDV 3A | Mouse | Clone 2C2 |

For live experiments, PK-15 G3BP1-RFP cells were seeded into Nunc™ Lab-Tek™ II chambered cover glasses (Thermo Fisher). The day after, cells were infected and, after inoculation, incubated with 10 μg/ml Hoechst H3570 (Thermo Fisher) for 30 minutes, and imaged for up to 7 hours after infection in Leibovitz’s L-15 Medium (Thermo Fisher) supplemented with 1 % FBS. Cells were visualized and movies were made with Leica SP8 CLSM inverted microscope using a 40X objective. Confocal images were analysed utilizing Image J by following the assembly and disassembly of G3BP1-foci in cells that develop CPE at around 6 hours of infection.

### Immunoblotting

Cells were seeded in 24-well plates to be around 80% confluent at the indicated times. Cells were treated as required, then media was removed, washed with cold PBS, lysed with 1x red Loading buffer supplemented with DTT (Cell Signalling) and boiled at 95 °C for 5 min. Lysates were separated in 4-20% gradient gel with 1x SDS running buffer (25 mM Tris, 192 mM glycine, and 0.1% SDS) and proteins were transferred to a nitrocellulose membrane using the Pierce Power Station and 1-step Transfer Buffer (Thermo Scientific), or by wet transfer using an appropriate buffer (25 mM Tris, 192 mM glycine, and 20% methanol). Membranes were blocked in 5% BSA in TBS-Tween (TBS-T) for 1 h at RT and then incubated with primary antibodies (Table **4**) overnight at 4 °C.

**Table 4.** Antibodies used on western blot staining.

| Primary antibodies used in WB | Host species | Reference |
| --- | --- | --- |
| p-STAT1 (Y701) | Rabbit | Cell Signalling Technology, #9167 |
| STAT1 | Mouse | Cell Signalling Technology, #9176 |
| p-IRF3 (S396) | Rabbit | Cell Signalling Technology, #4947 |
| IRF3 | Rabbit | Cell Signalling Technology, #8478 |
| IFIT2 | Rabbit | Protein Tech, 12604-1-AP |
| IRF1 | Rabbit | Cell Signalling Technology, #8784 |
| ISG15 | Rabbit | Protein Tech, 15981-1-AP |
| p-eIF2α (S52) | Rabbit | Cell Signalling Technology, #3597 |
| eIF2α | Rabbit | Cell Signalling Technology, #9722 |
| Puromycin | Mouse | Millipore, MABE343 |
| GAPDH | Mouse | Ambion, AM4300 |
| G3BP1 | Rabbit | Merck, G6046 |
| G3BP1 | Mouse | BD Transduction, #611127 |
| G3BP2 | Rabbit | Abcam, ab86135 |
| eIF4GI | Rabbit | Cell Signalling Technology, #2858 |
| FMDV 3A | Mouse | Clone 2C2 |

Next day, membranes were washed 3 times with TBS-T and subsequently incubated with secondary antibodies (Dako 1:5000, or LiCor IRDye 1:10,000) diluted in blocking buffer for 1 h at RT. After three washes with TBS-T, secondary antibodies were labelled with Clarity Western ECL Substrate (Bio-Rad) when required, and membranes were imaged using Bio-Rad or Odyssey Imaging systems. Band intensity was quantified and analysed using Image Studio Lite (Ver.5.2). Total protein was detected using ponceau S staining.

### Preparation of RNA samples and RT-qPCR

Cells were seeded in 24-well plates to be around 80% confluent at the indicated times. Cells were treated as needed, washed with cold PBS, and total RNA was extracted using a QiaGen RNAeasy kit (QiaGen, #74104) following the manufacturer’s instructions. Purified RNA was quantified on Nanodrop, and 500 ng of RNA was subject to reverse transcription using M-MLV Reverse Transcriptase (Invitrogen, #28025013). Next, real-time quantitative PCR (qPCR) was performed in duplicate with 25 ng of template, specific primers (Table **5**) used at a final concentration of 0.25 μM and PowerUp™ SYBR™ Green Master Mix (Applied Biosystems), following manufacturer’s instructions, using the Quant Studio 5 software (Applied Biosystems). The comparative Ct method was used; ΔCT values were obtained by normalizing the Ct values from genes of interest to *GAPDH*, and the 2-ΔCT method was used to calculate its relative expression in each sample.\

**Table 5.**
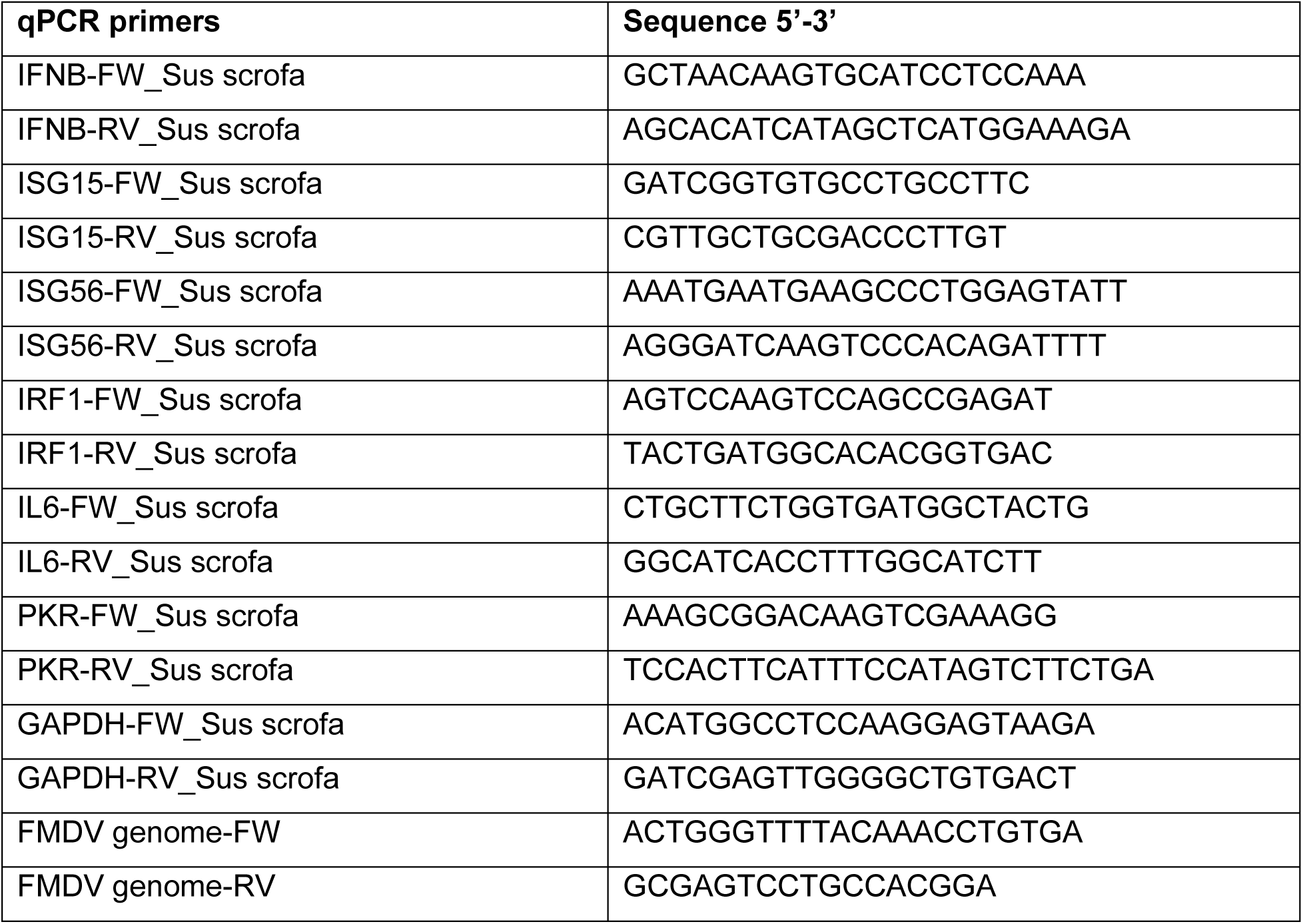
qPCR primers.

